# The neuroprotective effects of estrogen and estrogenic compounds in spinal cord injury

**DOI:** 10.1101/2022.10.30.514438

**Authors:** Artur Shvetcov, Marc J. Ruitenberg, Fabien Delerue, Wendy A. Gold, David A. Brown, Caitlin A. Finney

## Abstract

Spinal cord injury (SCI) occurs when the spinal cord is damaged from either a traumatic event or disease. SCI is characterised by multiple injury phases that affect the transmission of sensory and motor signals and lead to temporary or long-term functional deficits. There are few treatments for SCI. Estrogens and estrogenic compounds, however, may effectively mitigate the effects of SCI and therefore represent viable treatment options. This review systematically examines the pre-clinical literature on estrogen and estrogenic compound neuroprotection after SCI. Several estrogens were examined by the included studies: estrogen, estradiol benzoate, Premarin, isopsoralen, genistein, and selective estrogen receptor modulators. Across these pharmacotherapies, we find significant evidence that estrogens indeed offer protection against myriad pathophysiological effects of SCI and lead to improvements in functional outcomes, including locomotion. A STRING functional network analysis of proteins modulated by estrogen after SCI demonstrated that estrogen simultaneously upregulates known neuroprotective pathways, such as HIF-1, and downregulates pro-inflammatory pathways, including IL-17. These findings highlight the strong therapeutic potential of estrogen and estrogenic compounds after SCI.

## 1. Introduction

There is growing evidence that estrogen and estrogenic compounds are neuroprotective. Estrogens are a class of steroid hormones synthesized in endocrine organs as well as in the central nervous system (CNS) (Balthazart & Ball, 2006; Cui et al., 2013). Despite estrogen being conceptualized as a female hormone, males do synthesize some estrogen in the testes and other non-reproductive tissues (Cui et al., 2013). Further, males synthesize estrogen in the brain, with evidence suggesting little sex difference in the brain’s ability to locally synthesize estrogen (Cornill et al., 2012; Cui et al., 2013; Stoffel-Wagner et al., 1999). The predominant estrogen in the body is 17β-estradiol and, as such, the term estrogen hereafter refers to 17β-estradiol. Estrogens, and estrogenic compounds that mimic estrogen’s effects, bind to estrogen receptors (ERs) including the ERα, ERβ, and the G protein-coupled ER (GPER) (Paterni et al., 2014; Prossnitz & Barton, 2011). Recent studies have also identified novel ERs, such as ERX and an STX-sensitive Gq-membrane ER. These newer ERs, however, remain poorly characterized and have not been extensively studied (Srivastava et al., 2013). Upon binding to ERs, estrogen acts via both classical (genomic) and non-classical (non-genomic) mechanisms. Classical mechanisms involve the estrogen-estrogen receptor complex acting as a transcription factor by binding to estrogen response elements (EREs) on DNA sequences, or through the co-recruitment of other transcription factors. Non-classical mechanisms involve the rapid initiation of downstream signalling cascades and second messenger systems (Finney et al., 2020). Initial studies of estrogen-based hormone replacement therapies in women showed there was a decreased risk of developing neurodegenerative conditions, providing the initial indications that estrogen may be protective (Dubal, 2002). Subsequent studies have shown that estrogens are indeed protective against many neurodegenerative conditions including, but not limited to, multiple sclerosis (Spence & Voskuhl, 2012), Alzheimer’s disease (Simpkins & Singh, 2008), Parkinson’s disease (Morale et al., 2006), as well as traumatic brain and spinal cord injury (Chakrabarti et al., 2015). Although the specific mechanisms underlying estrogenic neuroprotection are largely unknown, estrogen has been shown to suppress pro-inflammatory pathways, oxidative stress, and apoptosis and simultaneously induce neurogenesis (Chakrabarti et al., 2015).

Estrogen and estrogenic compounds may effectively mitigate the effects of spinal cord injury (SCI). SCI occurs when the spinal cord is damaged from either a traumatic event or disease, affecting the transmission of sensory and motor signals and leading to either temporary or long-term changes (Ahuja et al., 2017; Marino et al., 2002). After SCI, patients typically experience a low quality of life characterized by a lack of mobility and a loss of independence, as well as an increased risk of mortality (Ahuja et al., 2017). Recent estimates suggest that the prevalent cases of SCI globally are between 25 and 30 million people, with incidence rates from 13 to 26 per 100,000 people and a an average cost of approximately $1.5 to 3 billion per annum, per country (Collie et al., 2010; GBD 2016 Traumatic Brain Injury and Spinal Cord Injury Collaborators, 2019; Krueger et al., 2013; McDaid et al., 2019). Despite the personal and economic burden of SCI, there are few treatments available. Another hormone, the glucocorticoid analogue methylprednisolone was considered a first line pharmacotherapy for SCI, however it is associated with adverse effects including gastric bleeding, and exacerbation of injury-related immunosuppression contributing to wound infection and a lower lymphocyte nadir. This led to the discontinuation of its use in most countries (McDonald & Sadowsky, 2002). Currently, the main treatment used for SCI is early surgical decompression with the critical window for this being within the first 24 hours after injury (Ahuja et al., 2020). Further complicating treatment is the nature of SCI, which is characterized by multiple distinct injury phases. The acute phase, occurring immediately after injury, consists of edema, hemorrhage, inflammation, and neuronal and myelin changes. This is followed by an intermediate phase characterized by gliosis and vascular changes that occurs across days to weeks after the initial injury and leads to significant secondary neurodegeneration. Finally, the late phases of SCI last months to years and are characterized by degeneration, glial and fibrous scars, and cavity formation (Ahuja et al., 2017; Norenburg et al., 2004).

The aim of the present systematic review is to (re-)evaluate if estrogen and estrogenic compounds may be neuroprotective against the effects of SCI. While there have been previous reviews on estrogen and neuroprotection in acquired central nervous system (CNS) injuries, they have typically combined research on SCI with other neurodegenerative injuries like traumatic brain injury. Although looking at estrogen’s neuroprotective role across injuries and conditions is valuable, it fails to consider differences in the nature of the injury and subsequent pathophysiological sequelae, such as the immune dysfunction seen in SCI (Allison & Ditor, 2014). Previous reviews have also not considered a role for estrogenic compounds in SCI. This review therefore marks the first attempt at systematically examining the literature on estrogen and estrogenic compounds in relation to neuroprotection after SCI, offering a comprehensive analysis of this literature base.

## 2. Methods

### 2.1. Literature search

Three databases were used to conduct this systematic review: PubMed, Scopus, and Web of Science. Independent searches of each database were performed on October 17, 2022. The search strategy used included the following key terms: estrogen, estradiol, spinal cord injury and were limited to the title and abstract. Results from all three searches were combined and duplicates removed. The resultant titles, abstracts, and full texts were rated based on eligibility criteria developed *a priori* (Table 1).

**Table 1.** Selection criteria

|  |
| --- |
| **Inclusion Criteria** |
| Male or female subjects and/or motoneuron cell culture |
| Direct spinal cord injury or spinal cord injury animal model |
| Administration of estrogen or estrogenic compound(s) |
| Spinal cord-specific and/or functional outcomes described |
| Studies written in English |
| Peer-reviewed article |
| **Exclusion Criteria** |
| No estrogen or estrogenic compounds used |
| Use of estrogen or estrogenic compound as an adjunctive therapy |
| No spinal cord injury |
| Did not examine either spinal cord pathology or functional outcomes |
| Not in English |
| Not a primary study |

### 2.2. Eligibility criteria

The goal of the present review was to establish if estrogen and estrogenic compounds have a neuroprotective role against the pathophysiological effects of SCI. The citations obtained from the literature searches were screened based on (a) male or female subjects and/or use of motoneuron cultures, (b) direct spinal cord injury or spinal cord injury animal model, for *in vitro* studies, (c) administration of estrogen or estrogenic compound(s), and (d) spinal cord-specific and/or functional outcomes described. Included studies were limited to those written in English and published in peer-reviewed journals (Table 1). Studies were excluded on the basis of (a) no estrogen or estrogenic compound used (e.g. use of the natural estrus cycle), (b) use of estrogen or estrogenic compound as an an adjunctive therapy (e.g. estrogen combined with stem cell therapy), (c) no spinal cord injury or model, (d) did not examine spinal cord or functional outcomes (e.g. brain outcomes), (e) not in English, and/or (f) not a primary study (e.g. a review) (Table 1). Importantly, the titles and abstracts were screened liberally, and exclusion criteria were not strictly applied until the full text was read. Titles and abstracts are often ambiguous with respect to the methods used and postponing the strict application of the exclusion criteria until the full text stage ensured that no studies were incorrectly excluded.

### 2.3. Protein-protein interaction and functional network analysis

Almost all of the included studies looked at the neuroprotective effects of estrogen and estrogenic compounds on molecular pathophysiology after SCI including gene expression and protein changes. Importantly, SCI recruit proteins that exist in functional networks where they interact to lead to a final outcome (Sevimoglu & Arga, 2014). To date, no studies have used high throughput analyses after estrogen or estrogen compound treatment, thus limiting our current understanding of the networks and pathways that may underlie neuroprotection. To begin to better understand these networks and pathways, we performed a protein-protein interaction and functional network analysis in STRING v11 (Szklarczyk et al., 2019). Proteins that were reported to be significantly affected by estrogen or estrogenic compounds (*p* < 0.05), either by modulating synthesis or phosphorylation, were included in the analysis (Finney et al., 2020). Importantly, proteins were only included if at least half of the studies reported a finding in the same direction. For example, if seven studies looked at levels of the protein caspase-3, at least four of those would have to show that estrogen decreased caspase-3 levels to be included. Functional connectivity between proteins was determined using the protein-protein interaction (PPI) enrichment value (Szklarczyk et al., 2019; Szklarczyk et al., 2016). Active interaction sources were limited to experiments, databases, co-expression, neighbourhood, and gene fusion. Interaction sources that resulted from text mining or co-occurrence were excluded. The minimum interaction score required was a medium confidence of 0.400. Proteins involved in functional or biological processes and their localization in the cell were identified using the Gene Ontology (GO) database (Gene Ontology Consortium, 2004). The contribution of the included proteins was also assessed using the Kyoto Encyclopedia of Genes and Genomes (KEGG) database (Kanehisa & Goto, 2000). To determine the significance of the functional and biological processes, cellular localizations, and pathways identified, the false discovery rate (FDR) statistic was used. Importantly, this statistic corrects for multiple comparisons and is commonly used in proteome data analytics (Rouam, 2013; Szklarczyk et al., 2019; Szklarczyk et al., 2016). This approach has proved to be valuable in previous work identifying potentially novel mechanisms underlying estrogenic regulation of memory (Finney et al., 2020).

## 3. Results

### 3.1. Literature search

Three databases were searched PubMed, Scopus, and Web of Science, yielding a total citation count of 777 citations. After duplicate citations were removed, a total of 493 citations remained. Titles and abstracts were then screened based on inclusion and exclusion criteria developed *a priori* (Table 1). A total of 88 full texts were reviewed for inclusion in the current systematic review. Of these, the most common cause of exclusion was due to no estrogen or estrogenic compound used (n = 12), not in English (n = 6), use of estrogen or estrogenic compound as an adjunctive therapy (n = 4), no spinal cord injury (n = 1), or not a primary study (n = 1) (Figure 1). In addition to the excluded full texts based on the *a priori* exclusion criteria, another three papers were excluded as they were unable to be located despite the best efforts of the authors. A further three studies examining estrogen treatment were also excluded due to significant methodological concerns, casting doubt on the results reported: these included identifying statistical significance where the reported averages and standard error of the means between the experimental and control groups were near identical; unclear doses of estrogen used in experimental groups; and/or a lack of statistical significance on any measures between the sham and SCI groups, indicating there was no specificity of the observed response.

**Figure 1.**
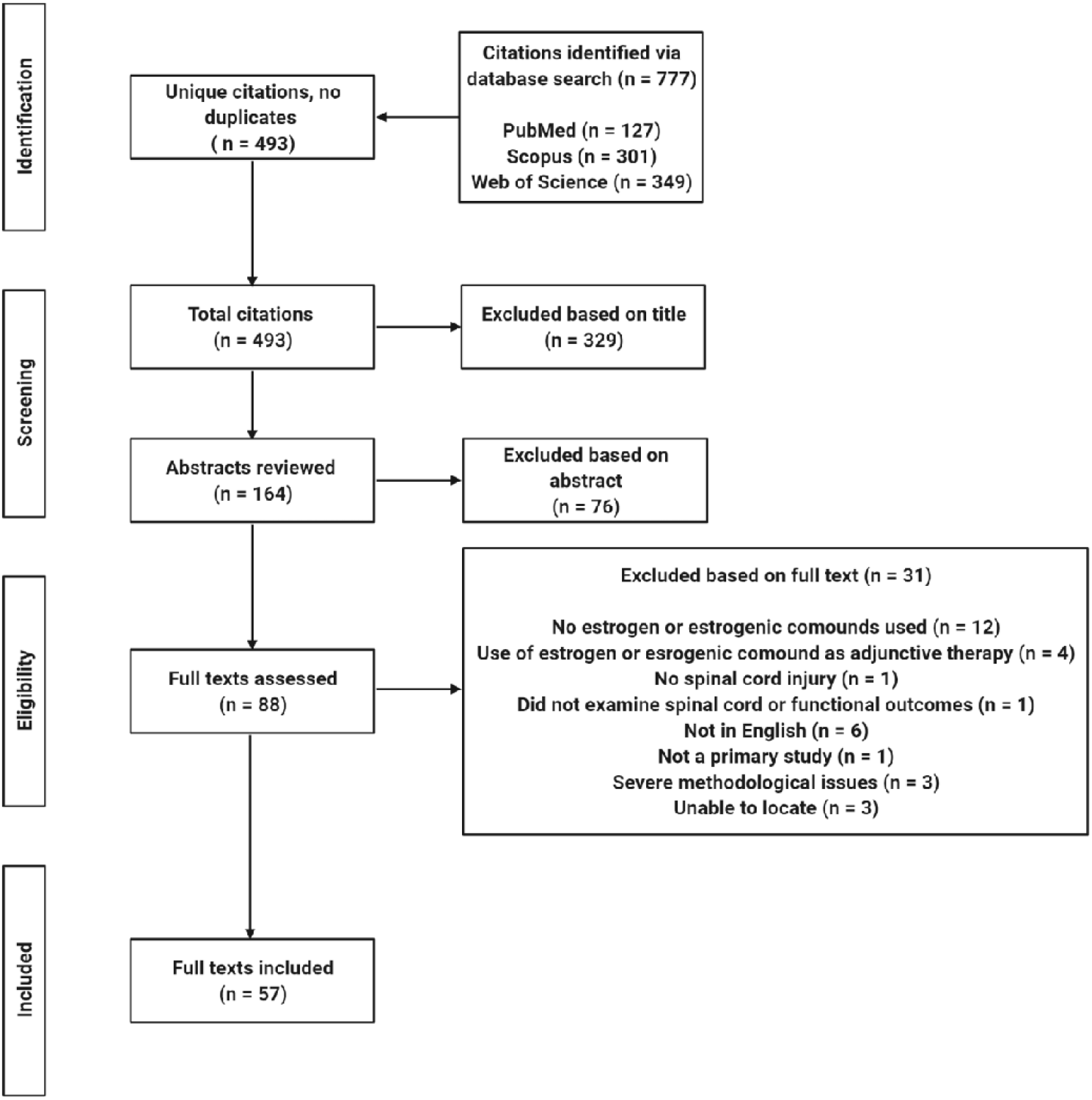
Literature database search inclusion and exclusion process.

### 3.2. Characteristics of included studies

As outlined in Table 2, almost all of the included studies in the present review used *in vivo* rodent models for SCI (n = 53/57) and only a few used cell culture (n = 5/57); one study used both cell culture and a rodent model of SCI and this was accounted for in each of the respective total n’s. Of those using rodent models, most used rats, either Sprague Dawley (n = 37) or Wistar (n = 10). Only six studies used mice including C57Bl/6J (n = 4), C57Bl/6J x 129S4/SvJae (n = 1), and CD1 (n = 1) (Table 2). The age of the animals was variable, with ages ranging from two months to one year. Most studies, however, did not specify the age of the animal, simply calling them either “adult” (n = 32) or “young adult” (n = 2) (Table 2). Importantly, previous research has documented the effects of age on outcomes after SCI, including, for example, neuropathic behaviour (Gaudet et al., 2021; Gwak et al., 2004), lesion pathology (Hooshmand et al., 2014), as well as microglial (Hooshmand et al., 2014) and inflammatory responses (Kumamaru et al., 2011). Lack of this information marks an important data gap in the literature and future studies should clearly identify the age of the animals used. Most studies (n = 41) used male rodents and only ten used females. There was a clear sex-specific effect of gonadal status of the animal, with half the studies using ovariectomized females and only one orchidectomizing the males (Table 2). Therefore, it is important that studies continue to focus on examining the neuroprotective effects of estrogen and estrogenic compounds in both intact and ovariectomized females to model SCI in both young (reproductive age) and older (menopausal) women.

**Table 2.** Characteristics of the subjects used in the included studies

| **Subject Characteristics** | **Number of Included Studies** |
| --- | --- |
| **Species and Strain** |  |
| Rat, Sprague Dawley | 37 |
| Rat, Wistar | 10 |
| Mouse, C57Bl/6J | 4 |
| Mouse, C57Bl/6J x 129S4/SvJae | 1 |
| Mouse, CD1 | 1 |
| Cell culture | 5 |
| **Age** |  |
| ~2 months old | 5 |
| ~3 months old | 5 |
| ~4 months old | 3 |
| ~5 months old | 1 |
| ~1 year old | 1 |
| Embryonic | 5 |
| Young adult | 2 |
| Adult | 32 |
| Not specified | 2 |
| **Sex** |  |
| Male | 41 |
| Female | 10 |
| Both | 2 |
| Embryonic (therefore not specified) | 5 |
| **Gonadal Status** |  |
| Intact | 48 |
| Ovariectomized | 5 |
| Orchidectomized | 1 |
| Embryonic (therefore not applicable) | 5 |

The included studies were highly heterogeneous with respect to the SCI models used. Most studies used contusion SCI (n = 36) via either a weight drop device or an impactor, although they used different size of weights. For brevity and consistency, the characteristics of the contusion SCI have been converted into g/cm or kdyn units to account for both weight drop and impactor SCI methods. The force of these contusion injuries was variable and ranged from a mere 10 g/cm to 40 g/cm for weight drops and 75-225 kilodynes (kdyn) for those using an impactor (Table 3). A further three studies did not include specific details about the nature of the injury in their respective papers, including either a lack of height of the weight drop, or no details at all. Only 11 studies used a compression injury, with the range of forces used variable from 15g to 55g (Table 3). There were also four studies that did not specify the force used, and simply stated that forceps were used for 30 seconds. One study also did not include any specific methodological details (Table 3). The remaining studies used either transection (n = 2) or electrolytic (n = 4) SCI (Table 3). There were no *in vivo* studies that used distraction, dislocation, or chemical SCI models. The included studies that used cell culture all employed a different insult to model SCI, with no overlap between them (Table 3). Whilst there was a range in the spinal cord vertebrae and segments undergoing SCI in the included papers, almost all focused on areas of the thoracic spine, with the vast majority targeting between T8-T10 (Figure 2). As such, there is need for studies to examine estrogenic neuroprotection in SCIs sustained to other areas of the spine.

**Table 3.** Characteristics of the SCI models used in the included studies

| SCI Characteristics | Number of Included Studies |
| --- | --- |
| **Contusion SCI (total n = 36 studies)** |  |
| 10 g/cm | 1 |
| 12.5 g/cm | 8 |
| 25 g/cm | 7 |
| 32 g/cm | 2 |
| 40 g/cm | 9 |
| 75 kdyn | 2 |
| 150 kdyn | 2 |
| 200 kdyn | 1 |
| 225 kdyn | 1 |
| 35g (no height specified) | 1 |
| 50g (no height specified) | 1 |
| Not specified | 1 |
| **Compression SCI (total n = 11 studies)** |  |
| Vascular clip, 15g, 1 min | 1 |
| Microvessel clip, 20g, 15 sec | 1 |
| Aneurysm clip, 24g (no time specified) | 1 |
| Aneurysm clip, 55g, 1min | 1 |
| Aneurysm clip, 1.37-1.72N, 1 min | 2 |
| Forceps (no force specified), 3sec | 3 |
| Forceps (no force specified), 20sec | 1 |
| Not specified | 1 |
| **Transection SCI (total n = 2 studies)** |  |
| Partial via projectile (1x1x10mm) | 1 |
| Complete via dissection scissors | 1 |
| **Electrolytic SCI (total n = 4 studies)** |  |
| 300μA for 90sec | 4 |
| **Cell Culture Chemical SCI (total n = 5 studies)** |  |
| Calcium ionophore | 1 |
| IFN-γ | 1 |
| Microglial cytokine | 1 |
| Oxygen-glucose deprivation | 1 |
| TNF-α toxicity | 1 |

**Figure 2.**
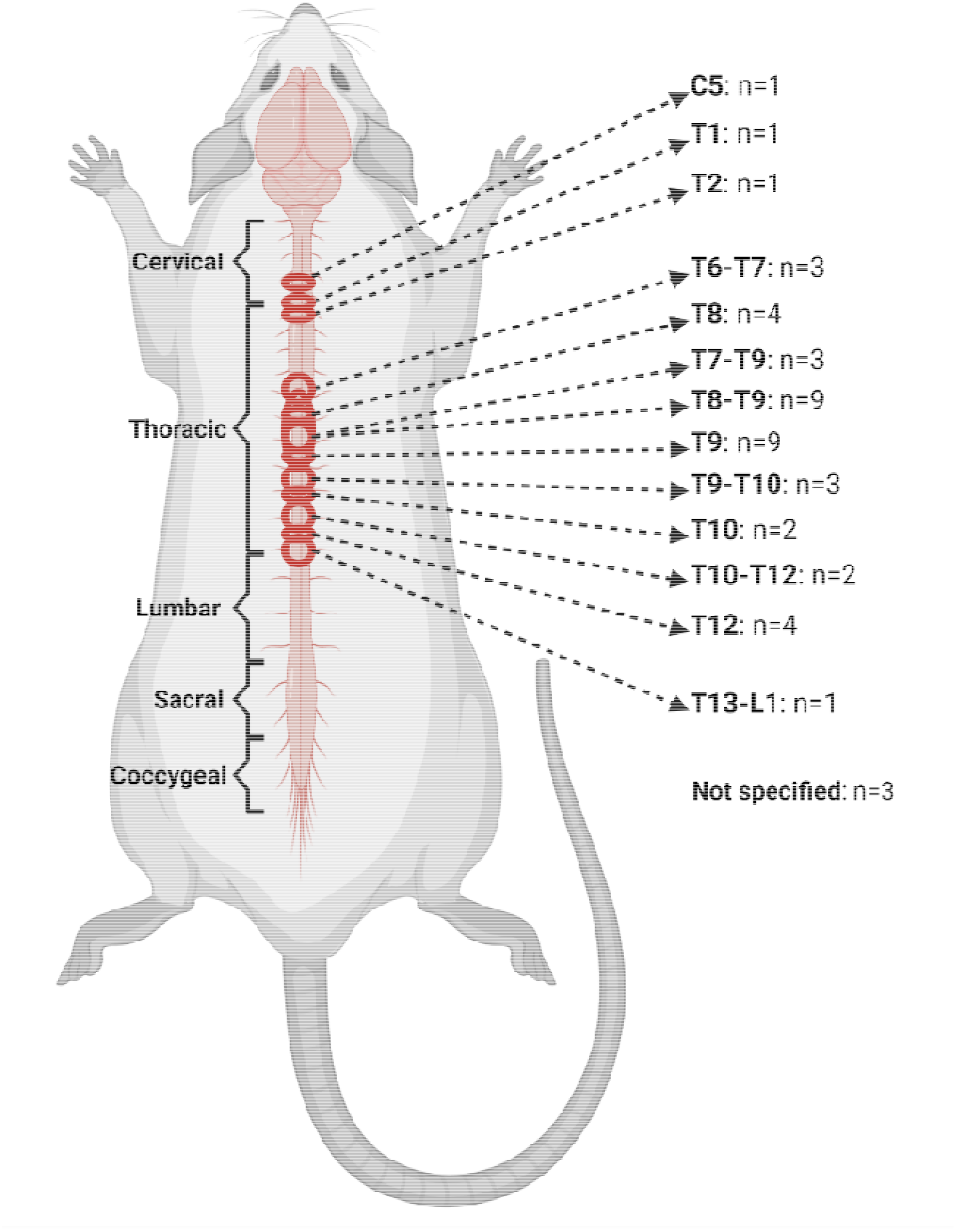
Location of SCI along rodent spinal cord used in the included studies. Includes n’s for the number of included studies targeting that area of the spinal cord. Schematic diagram of rodent spinal cord is not to scale, and spinal cord segment locations are approximate.

### 3.3. Estrogen protects against functional deficits after SCI

Overall, the studies included in the present review clearly demonstrate that estrogen, and its compounds, are protective against myriad functional outcomes after SCI (Table 4). Estrogen administered before, after, or both before and after SCI were consistently found to improve locomotion across a large range of doses from a mere 20pg/ml to 10g/kg and across a variety of locomotor measures. Estrogen improved the score on the Basso, Beattie, Bresnahan (BBB) locomotor rating scale. Here, effects tended to be limited to doses of estrogen that were at 0.08mg/kg and above (Table 4). Studies demonstrated that this improvement could be seen from as little as 72 hours after injury, with the effects lasting up to 42 days (Chaovipoch et al., 2005; Chen et al., 2015; Hu et al., 2012; Kachadroka et al., 2010; Kim et al., 2021; Lee et al., 2012; Letaif et al., 2015; Lin et al., 2016; Majidpoor et al., 2020; Majidpoor et al., 2021; Mosquera et al., 2014; Ni et al., 2018; Ritz & Hausmann, 2008; Samantaray et al., 2016; Siriphorn et al., 2012; Sribnick et al., 2005; Tan et al., 2020; Yune et al., 2004); one study found that long-term BBB score improvement could be seen irrespective of whether estrogen was given pre- or post-SCI (Yune et al., 2004). Estrogen’s effects on the BBB score were also seen irrespective of age, as one study compared two-month-old to one-year-old rats and found that estrogen improved BBB score in both age groups (Chaovipoch et al., 2005). It is likely, however, that there is a critical dosing window for estrogen, with one study finding that estrogen only improved BBB score if given between one to two hours after SCI but not at six or 12 hours after (Lee et al., 2012). Additionally, it appears that exogenous, but not endogenous, estrogen is critical for neuroprotection after SCI. One study demonstrated the aromatase inhibitor letrozole in female rats did not affect performance on the open field test, although it did reportedly have some effect on progressively decreasing the paw withdrawal threshold (Ghorbanpoor et al., 2014). In males, BBB score improvement was unaffected by orchidectomy (Kachadroka et al., 2010). There may be a sex effect; however, with one study showing that estrogen administered to ovariectomized females improved BBB score but not when administered to intact male rats (Swartz et al., 2007). It is worth noting that this effect was only seen on day 7 post-SCI and not on days 14 or 21. Further, other studies did find that estrogen improved BBB score in both ovariectomized (Lin et al., 2016; Mosquera et al., 2014) and intact (Cheng et al., 2016; Kim et al., 2021) female rats. Estrogen also improved post-SCI performance in several other locomotor tests. These included the Basso Mouse Scale (BMS) (Cheng et al., 2016), the inclined plane test (Cheng et al., 2016), Vermicelli handling test (Siriphorn et al., 2012), horizonal ladder test (Siriphorn et al., 2012), as well as the narrow beam test, although only at a higher dose (Ritz & Hausmann, 2008). Despite estrogen being shown to improve performance across a variety of locomotor measures, only a temporary improvement was seen using the rotarod task on day 14 after injury but not by day 28 (Afhami et al., 2016). In one study, distance travelled in the open field task was also not affected by estrogen (Naderi et al., 2014) and another showed a brief improvement on days 7 and 14 but not by day 28 after injury (Afhami et al., 2016). There was also no effect of estrogen found on locomotion using the catwalk gait analysis (Siriphorn et al., 2012).

**Table 4.**
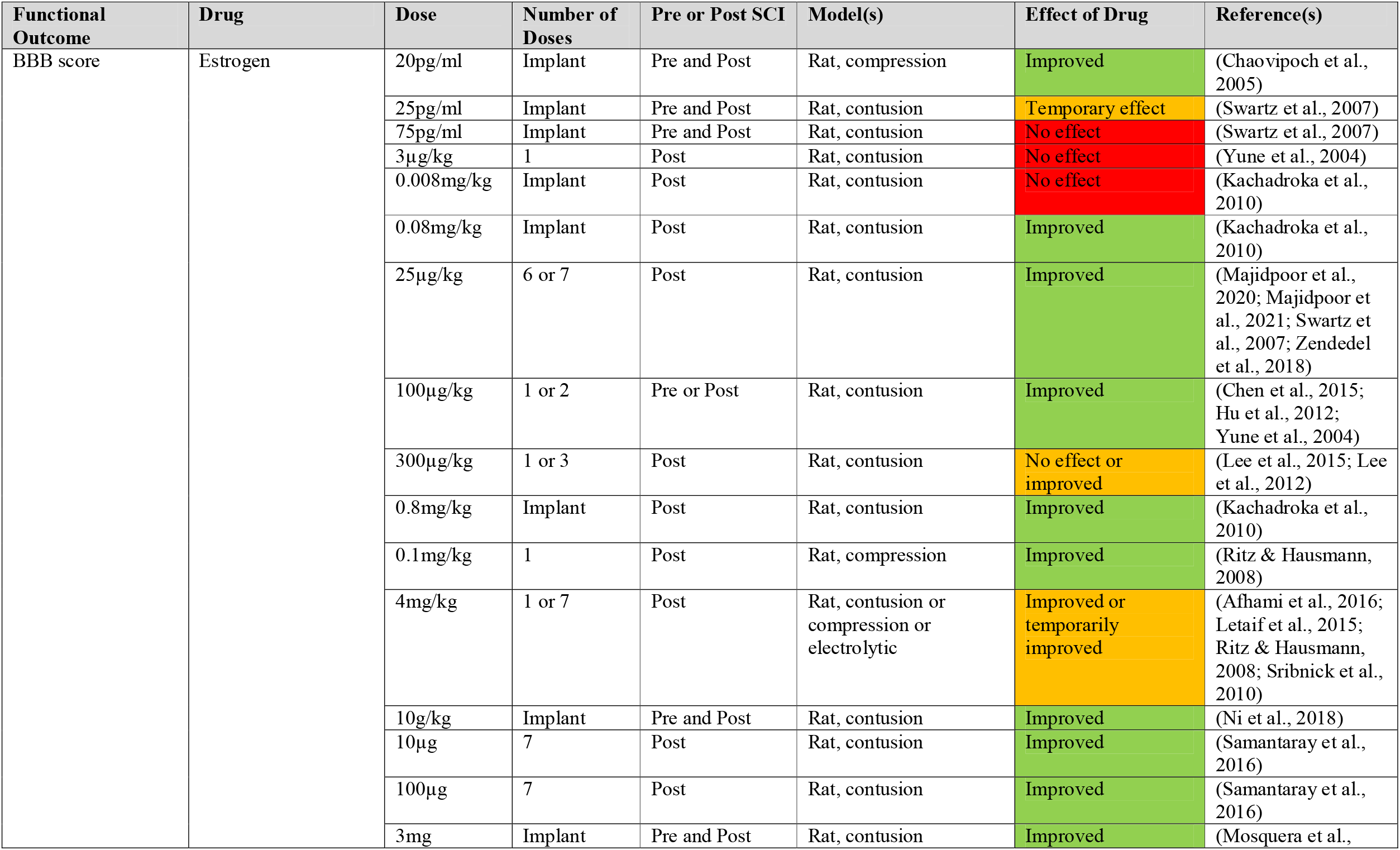
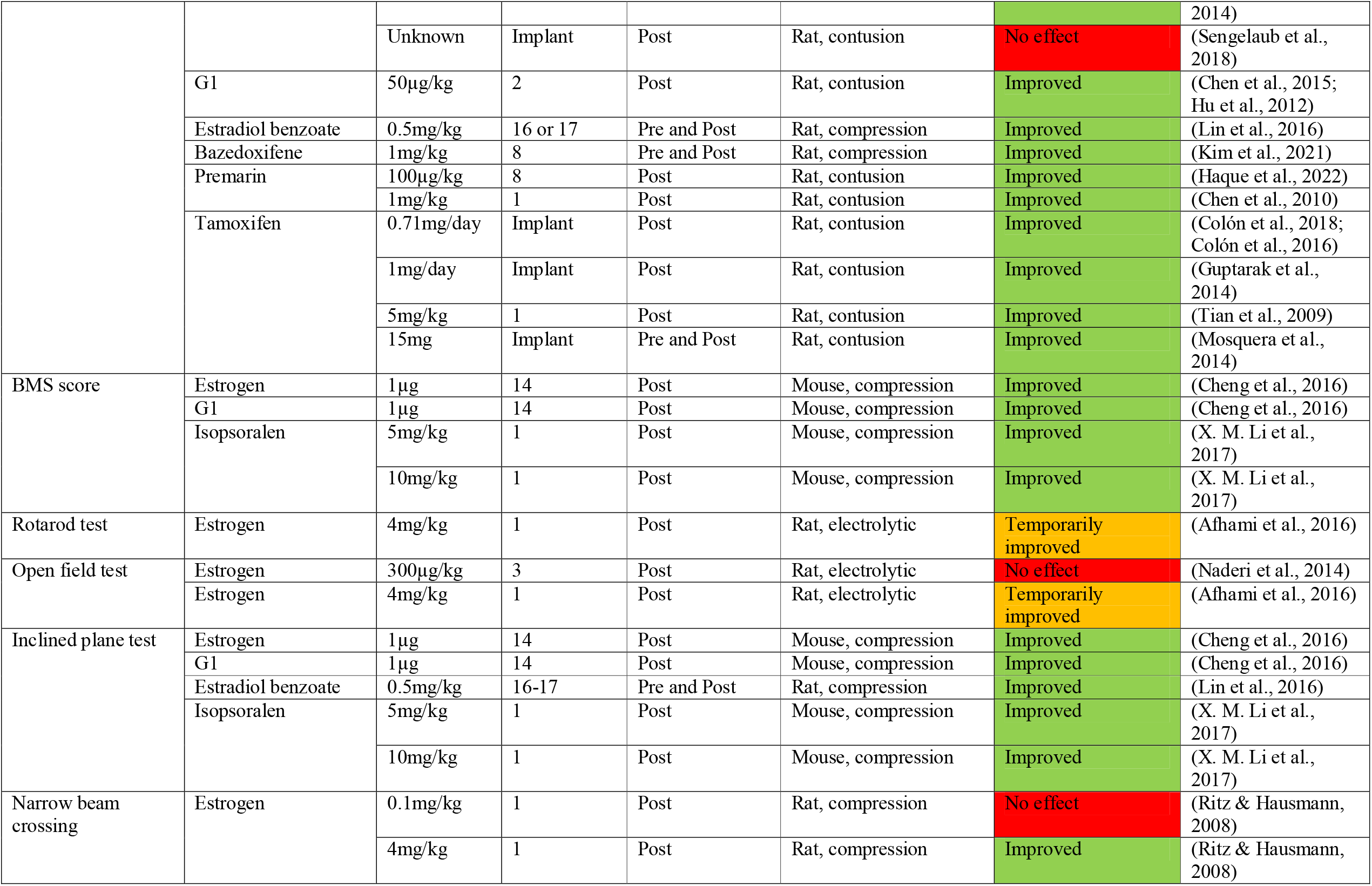
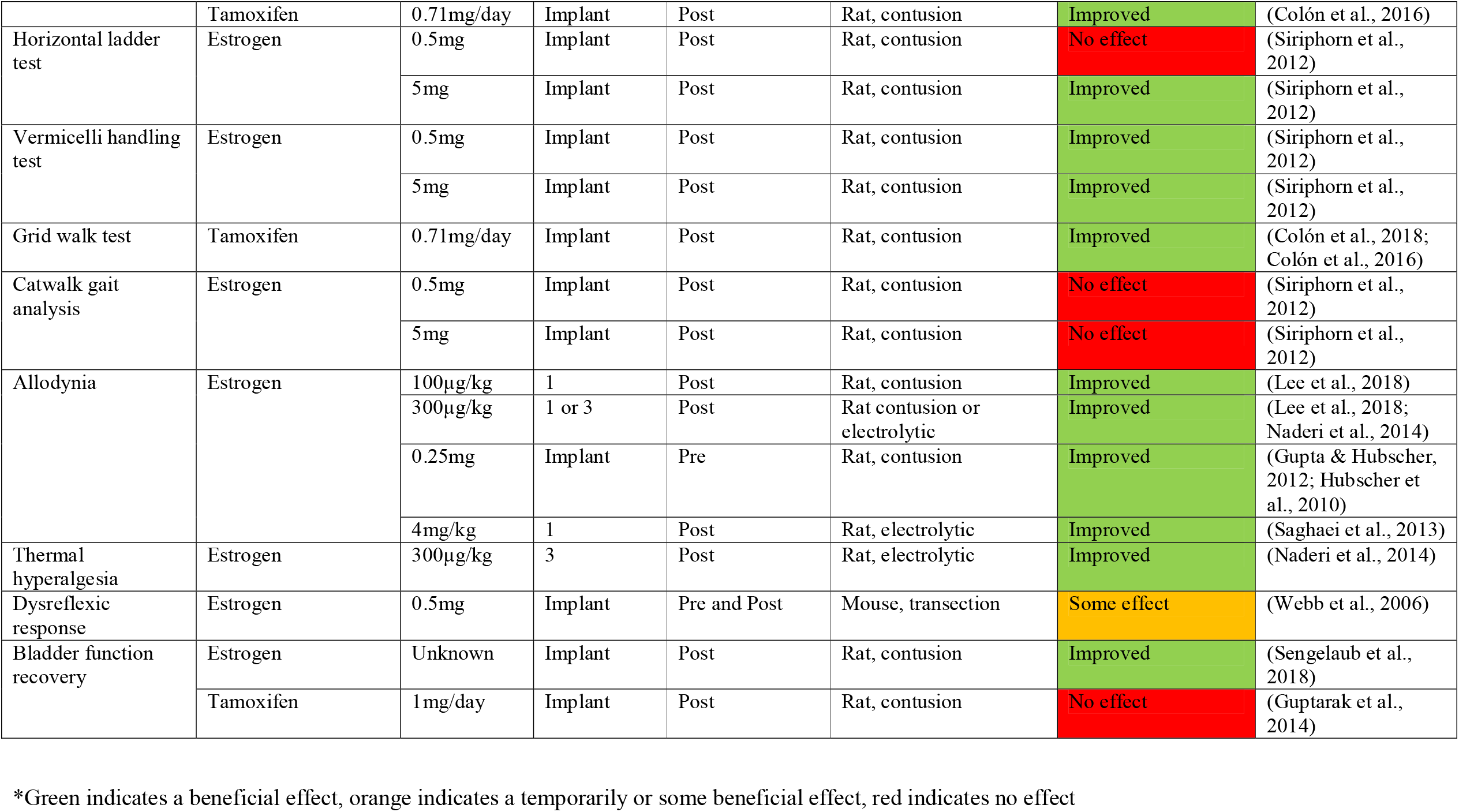
Effects of estrogen and estrogenic compounds on functional outcomes after SCI.

In addition to locomotion, several studies also looked at estrogen’s effects on other functional outcomes including allodynia, sensitivity to touch, thermal hyperalgesia, dysreflexic responses, and bladder function recovery. Estrogen improved allodynia across several different doses (Gupta & Hubscher, 2012; Lee et al., 2018; Naderi et al., 2014; Saghaei et al., 2013). It also decreased sensitivity to touch, which is increased in rodents after SCI (Hubscher et al., 2010). Thermal hyperalgesia (Naderi et al., 2014) as well as bladder function (Sengelaub et al., 2018) were also improved by estrogen administration (Naderi et al., 2014). With respect to dysreflexic responses, however, there was a differential effect of estrogen: it improved the dysreflexic response to colorectal distension but not to tail pinch (Webb et al., 2006).

### 3.4. Estrogenic compounds similarly protect against functional deficits after SCI

With respect to estrogenic compounds, our included studies examined five of them for neuroprotection: estradiol benzoate; tamoxifen; bazedoxifene; premarin; and isopsoralen (Table 4). Tamoxifen and bazedoxifene are first- and third-generation selective estrogen receptor modulators (SERMs), respectively (Komm et al., 2005).

Tamoxifen is approved for use in hormone receptor-positive breast cancers whereas bazedoxifene is used primarily to treat the effects of menopause, including osteoporosis. While estradiol benzoate, a synthetic estrogen, improved BBB score and performance on the inclined plane test in rats with compression SCI (Lin et al., 2016) tamoxifen and bazedoxifene both improved the neurological outcome from SCI, including BBB scores (Colón et al., 2018; Colón et al., 2016; Guptarak et al., 2014; Kim et al., 2021; Mosquera et al., 2014; Tian et al., 2009), narrow beam crossing (Colón et al., 2016), and grid walking (Colón et al., 2018; Colón et al., 2016). There was some evidence that tamoxifen has a critical time window for neuroprotection after SCI, as one study did demonstrate that tamoxifen only improved BBB score and performance on the grid walk task if given either immediately or six hours after SCI, but not 12 or 24 hours later (Colón et al., 2018). Another study, however, demonstrated that tamoxifen still improved BBB score and grid walking, irrespective of whether it was given either immediately or 24 hours post-insult, although only immediate (but not 24-hour) tamoxifen treatment improved narrow beam crossing (Colón et al., 2016). Despite these studies demonstrating a positive effect of tamoxifen on locomotion, it was not shown to have any beneficial effect on bladder function recovery after SCI (Guptarak et al., 2014).

Premarin, a synthetic estrogen used as a hormone replacement therapy (Fugh-Berman, 2010), was also found to improve BBB score after SCI (Chen et al., 2010; Haque et al., 2021). Here, however, it is very important to highlight that one of these studies (Haque et al., 2021) found that low dose Premarin (100µg/kg daily for seven days) caused adverse reactions in their male rats. Specifically, their urine contained a thick, dark blood that was accompanied by pale eyes and lethargy approximately five days into treatment; these adverse events resolved upon the cessation of Premarin therapy (Haque et al., 2021). No such side effects were reported for the other study where SCI recovery was also improved but where only a single high dose (1mg/kg) of Premarin was administered (Chen et al., 2010). It may be the case that Premarin should only be used acutely, and this again could be investigated by future studies. Finally, a traditional Chinese herbal medicine, isopsoralen, a naturally occurring phytoestrogen found in Psoralea corylifolia L. fruit, improved BMS score and performance on the inclined plane task after SCI in one study at both low- and high-dose treatment, presumably acting as a CNS SERM (X. M. Li et al., 2017; Wei et al., 2016).

### 3.5. The neuroprotective effects of estrogens on functional deficits after SCI are mediated by multiple ERs

As discussed above, estrogen and its compounds act by binding to ERs. Some studies have attempted to identify the specific ER subtypes involved via which estrogen improves the neurological outcome from SCI. To show receptor specificity, most studies used estrogen in combination with the ERα and ERβ antagonist ICI 182,780 (Howell et al., 2000). These studies demonstrated that estrogen’s effects on BBB score improvement (Lee et al., 2015; Lee et al., 2012), and performances on the Vermicelli handling and horizontal ladder tests (Siriphorn et al., 2012) were reduced by co-administration of ICI 182,780. The beneficial effect of estrogen on paw withdrawal after SCI was also decreased by ICI 182,780 (Lee et al., 2018). Another study, however, found that ICI 182,780 was unable to block the effect of estrogen on BBB score (Hu et al., 2012). Interestingly, although ICI 182,780 is an antagonist at ERα and β, it is also a documented agonist of the G protein-coupled estrogen receptor (GPER) (Meyer et al., 2010), suggesting that some positive benefits of estrogen may mediated by this receptor. In line with this idea, the same study found that, unlike ICI 182,780, a GPER1 antisense oligonucleotide was able to effectively block estrogen’s improvement in BBB score after SCI (Hu et al., 2012). A positive role for GPER1 was corroborated in another study, which used the GPER agonist G-1 and demonstrated that this also improved functional scores (Cheng et al., 2016). This study also found that administration of G-15, a high affinity GPER antagonist, blocked estrogen-induced improvement on BMS score after SCI in mice (Cheng et al., 2016). However, another study failed to demonstrated that G-15 affected estrogen’s effect on BMS in a rat model of SCI (Chen et al., 2015). This may account for these differences and should be investigated by future studies of injury severity in any such studies between mice and rats should also be considered. Only one study has attempted to investigate the contribution of estrogen effects of ERα verses ERβ, demonstrating that the selective ERα antagonist MPP dihydrochloride was able to block the effect of estrogen on BBB score in rat SCI (Mosquera et al., 2014).

### 3.6. Both estrogen and estrogenic compounds attenuate the (histo)pathological effects of SCI

Several of the included studies examined the effects of estrogen and estrogenic compounds on injury size and other histopathological characteristics after SCI (Table 5). Generally, estrogen, Premarin, and tamoxifen appeared to reduce the lesion size after moderate contusion (via weight drop or impactor) and compression SCI (Chen et al., 2010; Haque et al., 2021; Mosquera et al., 2014; Ni et al., 2018; Ritz & Hausmann, 2008; Yune et al., 2004) and increase the amount of tissue spared (Ni et al., 2018; Samantaray et al., 2016; Sengelaub et al., 2018; Sribnick et al., 2010). Further, estradiol benzoate and bazedoxifene decreased the necrotic tissue area (Kim et al., 2021; Lin et al., 2016) and the phytoestrogen isopsoralen decreased cavitation volume (X. M. Li et al., 2017). The neuroprotective effects of estrogen tended to be at the higher doses, with two lower dose studies finding no effect (Ritz & Hausmann, 2008; Swartz et al., 2007). Only one study showed that estrogen did not affect the lesion size using hematoxylin and eosin staining, however they noted that their protocol was unlikely to be sensitive enough to detect changes in lesion size in their moderate SCI lesion model (Letaif et al., 2015). One study also found that tamoxifen had no effect on lesion size (Pukos & McTigue, 2020). This study, however, reportedly used a very high (300mg/kg) dose of tamoxifen in mice. If correct, the human equivalent dose (HED) would be 3,700mg/kg (Reagan-Shaw et al., 2007), well outside of the highest daily dose of tamoxifen (120mg) used in humans making it hard to interpret these data (Chow et al., 2003). The effects of estrogen were demonstrated to be mediated by both ERα and GPER, using the antagonists MPP and G15 as well as the agonist G1 (Cheng et al., 2016; Mosquera et al., 2014).

**Table 5.**
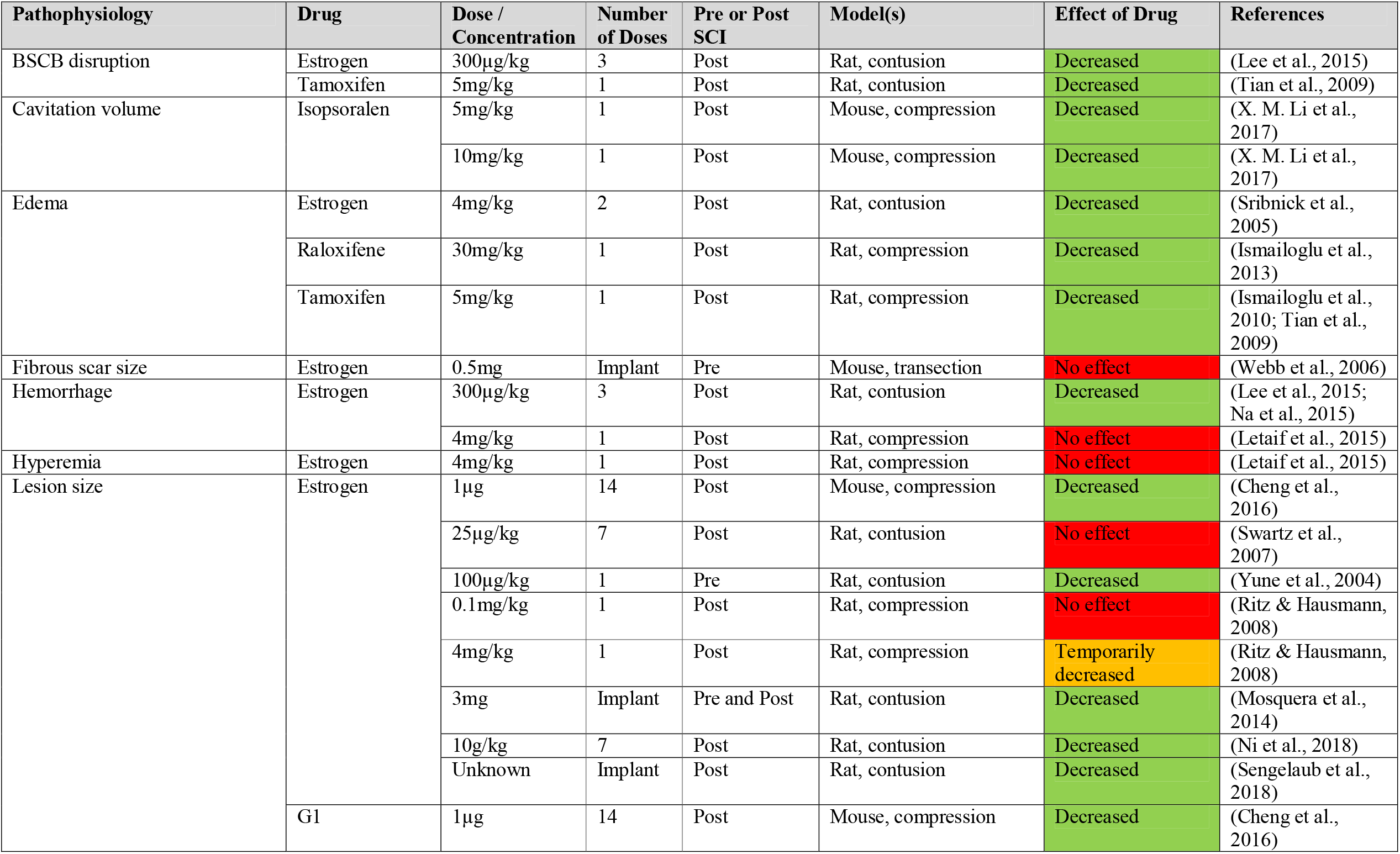
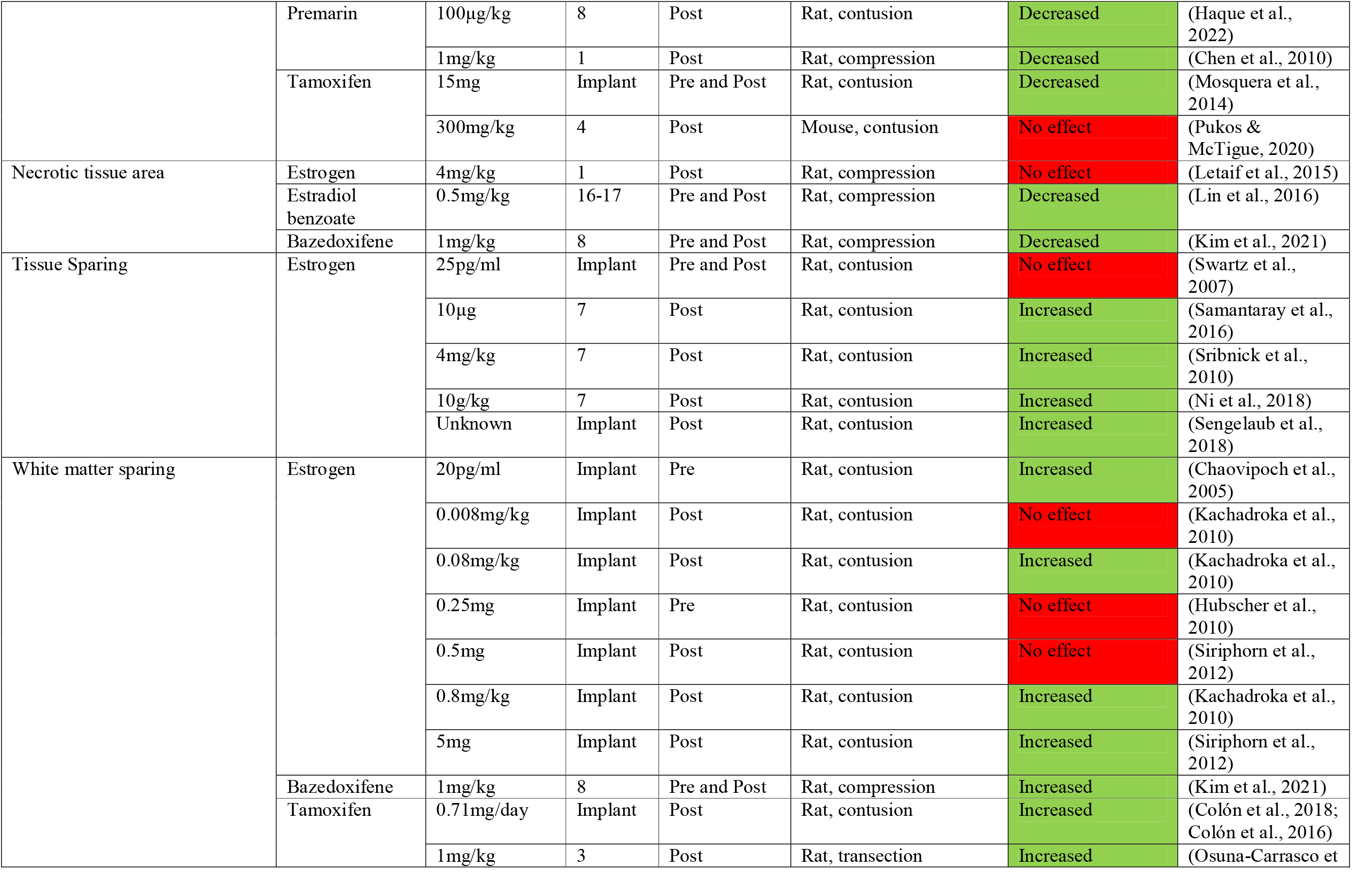
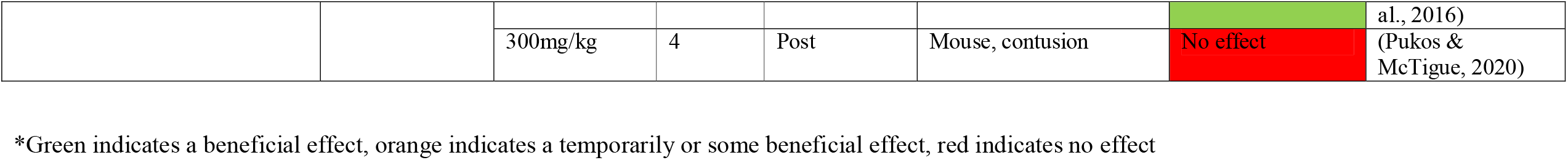
Effects of estrogen and estrogenic compounds on the (histo)pathological effects of SCI.

There were mixed effects of estrogens on white matter sparing (Table 5). Several studies showed that estrogen, tamoxifen, and bazedoxifene did increase the amount of white matter spared after SCI (Chaovipoch et al., 2005; Colón et al., 2018; Colón et al., 2016; Kachadroka et al., 2010; Kim et al., 2021; Osuna-Carrasco et al., 2016; Siriphorn et al., 2012). Other studies, however, found no effect (Hubscher et al., 2010; Kachadroka et al., 2010; Pukos & McTigue, 2020; Siriphorn et al., 2012). These differences were independent of the SCI model, as the studies included a mix of crush injuries, hemicontusion, and moderate contusion. Further, neither age (Chaovipoch et al., 2005) nor gonadal status (Kachadroka et al., 2010) were shown to affect estrogen’s ability to spare white matter. Interestingly, one study found that the non-specific ERα and β inhibitor ICI 182,780 was unable to affect estrogen’s efficacy (Siriphorn et al., 2012). This suggests that estrogen’s ability to spare white matter may indeed be more dependent on GPER and/or possibly involve other non-ER-dependent mechanisms. In cancers, for example, estrogen has been shown to exert ER-independent effects via its metabolites (Yue et al., 2013). This may account for the disparate findings of studies looking at white matter sparing after estrogen administration and warrants follow up study.

With respect to oedema and blood spinal cord barrier (BSCB) after SCI, estrogens showed a beneficial effect (Table 5). Estrogen, raloxifene, and tamoxifen were all shown to decrease odema and reduce the disruption to the BSCB (Ismailoglu et al., 2010; Ismailoglu et al., 2013; Lee et al., 2015; Sribnick et al., 2005; Tian et al., 2009). There was also a critical time window for estrogen neuroprotection against oedema and BSCB disruption. The positive effects were only seen if estrogen was given 1-2 hours after injury but not six hours later (Lee et al., 2015; Sribnick et al.,

### 3.7. Estrogen and estrogenic compounds reduce cell death, improve cell viability, and increase the number of remaining cells after SCI

Both estrogen and estrogenic compounds were consistently shown to have neuroprotective effects on cells after SCI (Table 6). Specifically, estrogen, estradiol benzoate, Premarin, bazedoxifene, tamoxifen, and genistein, decreased apoptosis, improved cell viability, and increased the number of remaining neurons after SCI (Chaovipoch et al., 2005; Chen et al., 2015; Chen et al., 2010; Colón et al., 2018; Das et al., 2011; Kachadroka et al., 2010; Kim et al., 2021; Lin et al., 2016; McDowell et al., 2011; Siriphorn et al., 2012; Smith et al., 2009; Sribnick et al., 2010).

**Table 6.**
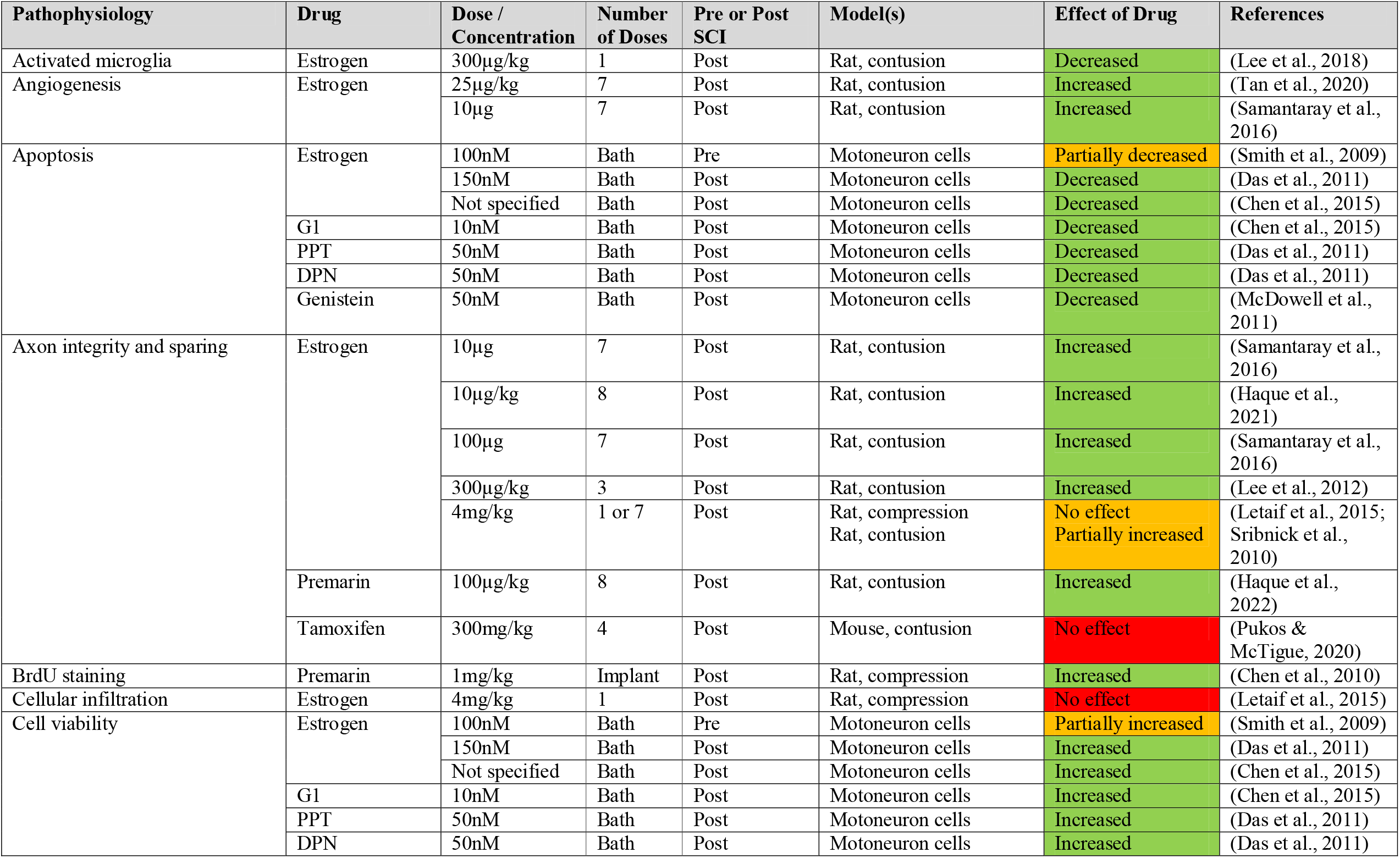
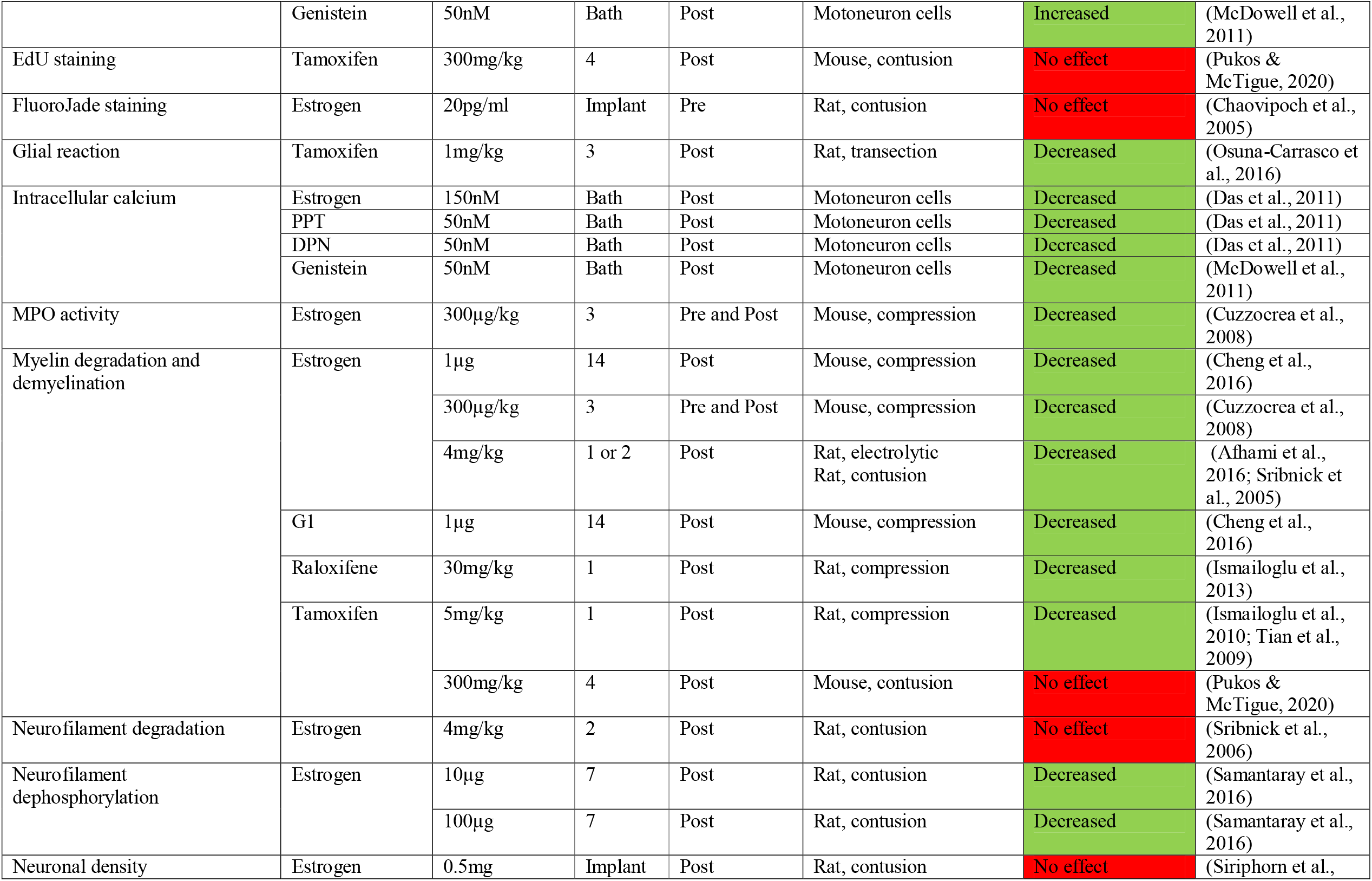
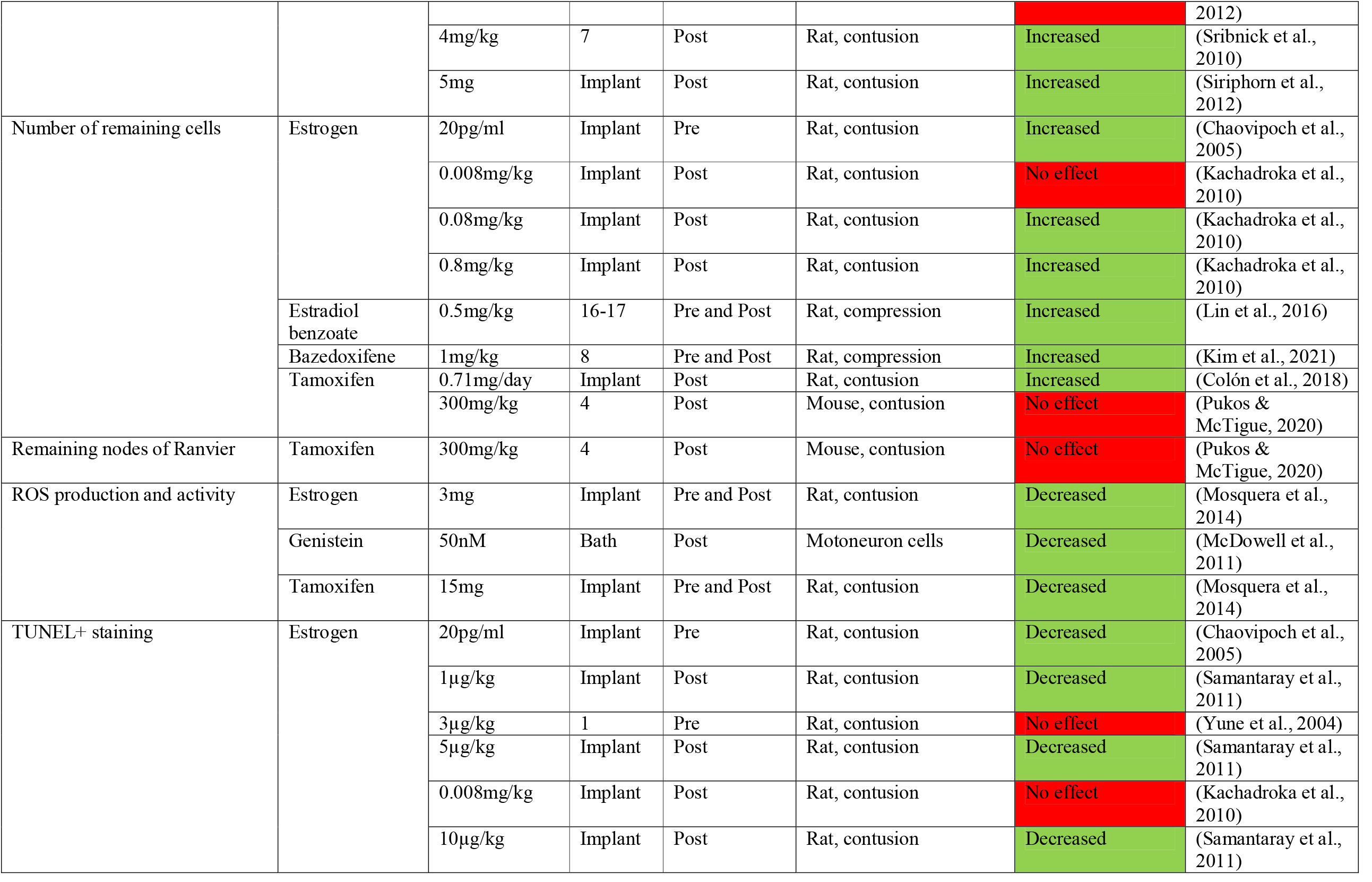
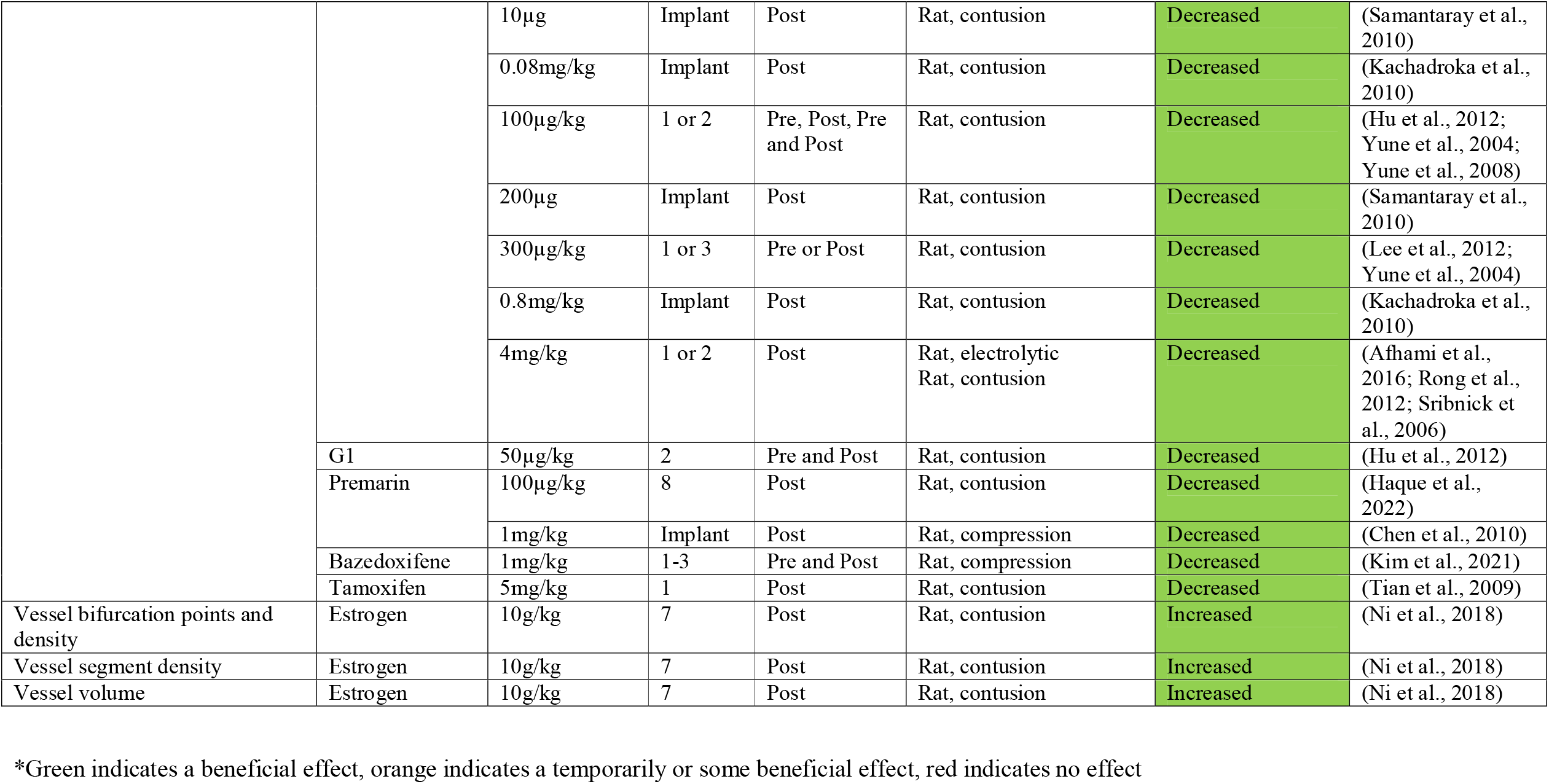
Effects of estrogen and estrogenic compounds on cellular pathophysiology after SCI

In line with this, numerous studies demonstrated that estrogen, Premarin, bazedoxifene, and tamoxifen decreased TUNEL positive staining, a marker of DNA fragmentation during the final stages of apoptosis (Afhami et al., 2016; Chaovipoch et al., 2005; Chen et al., 2010; Haque et al., 2021; Hu et al., 2012; Kachadroka et al., 2010; Kim et al., 2021; Lee et al., 2012; Rong et al., 2012; Samantaray et al., 2010; Samantaray et al., 2011; Sribnick et al., 2006; Tian et al., 2009; Yune et al., 2004; Yune et al., 2008). The beneficial effect of estrogen on decreased TUNEL-positive cells was independent of age, with both two-month and one-year old rats demonstrating estrogen-mediated decreases (Chaovipoch et al., 2005). Gonadal status, however, did influence estrogen’s ability to decrease TUNEL-positive cells. One study found intact males have a better response compared to orchidectomized males (Kachadroka et al., 2010). Estrogen’s effects on apoptosis were shown to be mediated by multiple ERs. Studies showed that administering propyl-pyrazole-triol (PPT), diarylpropionitrile (DPN), and G1 all mimicked estrogen’s ability to decrease apoptosis after SCI (Das et al., 2011). Further confirming this, studies blocking the GPER, using G15 or an antisense oligonucleotide knockdown, and ERα and β, using ICI 182,870, prevent estrogen-mediated decreases in apoptosis and improved cell viability (Chen et al., 2015; Das et al., 2011; Hu et al., 2012; Lee et al., 2012). Only one study found that estrogen-mediated protection against apoptosis after SCI was not affected by ICI 182,870 administration (Hu et al., 2012).

Estrogens also had beneficial effects on SCI-induced changes in cell morphology. Specifically, estrogen and Premarin were both shown to improve axon integrity and sparing after SCI (Haque et al., 2022; Haque et al., 2021; Lee et al., 2012; Samantaray et al., 2016; Sribnick et al., 2010). Two studies, however, demonstrated no effect. One study showed that estrogen (Letaif et al., 2015) and another, using a dose of tamoxifen well outside of the clinical range, showed that tamoxifen (Pukos & McTigue, 2020) had no effect on axon integrity and sparing.

### 3.8. Estrogen and estrogenic compounds prevent myelin degeneration and degradation after SCI

Several studies also examined the effects of estrogen and estrogenic compounds on myelin degradation and degeneration after SCI (Table 6). Estrogen, raloxifene, and tamoxifen were all shown to decrease the effects of SCI on myelin (Afhami et al., 2016; Cheng et al., 2016; Cuzzocrea et al., 2008; Ismailoglu et al., 2010; Ismailoglu et al., 2013; Sribnick et al., 2005; Tian et al., 2009). It was also shown that these positive effects of estrogens were mediated by ERα and β (Cuzzocrea et al., 2008) as well as GPER (Cheng et al., 2016). There was no effect, however, of estrogen on neurofilament degradation (Sribnick et al., 2006), although it was reported to protect neurofilament from dephosphorylation (Samantaray et al., 2016).

### 3.9. Estrogen and estrogenic compounds protect against SCI-induced intracellular calcium influx and ROS production and activity

Multiple estrogens were shown to protect against SCI-induced reactive oxygen species (ROS) production and activity including estrogen (Cuzzocrea et al., 2008; Mosquera et al., 2014), tamoxifen (Mosquera et al., 2014), and genistein (McDowell et al., 2011). Further, both estrogen and genistein decreased intracellular calcium influx after SCI (Das et al., 2011; McDowell et al., 2011). Only one study examined the role of ERs in these aspects of neuroprotection and showed that genistein’s effects could be blocked by the ERα and β antagonist ICI 182,870 (McDowell et al., 2011) (Table 6).

### 3.10. Estrogen and estrogenic compounds activate and recruit several established neuroprotective pathways after SCI

Across all of the included studies, estrogen and estrogenic compounds had myriad neuroprotective benefits, enforcing changes in RNA expression, protein levels, and phosphorylation status after SCI (Supplementary Table 1). Specifically, these changes led to beneficial modulation of cell proliferation, survival, differentiation, development, angiogenesis, and transcription pathways. Supplementary Table 1 outlines an exhaustive list of all these different processes as well as the transcripts and proteins that were shown to be involved in, and/or modulated by, estrogen and estrogenic compounds. Similar to the neuroprotective benefits above, these effects were mediated by multiple ERs including GPER (Chen et al., 2015; Cheng et al., 2016) and ERα and β (Cuzzocrea et al., 2008; Das et al., 2011; Haque et al., 2022; Lee et al., 2015; Lee et al., 2012; Siriphorn et al., 2012).

It is important to consider that proteins exist in functional networks that interact to generate a particular outcome (Sevimoglu & Arga, 2014). To date, however, no studies have used high-throughput analyses to examine the protein networks that underlie the effects of estrogen treatment after SCI. As such, there is currently a limited understanding of the networks, pathways, and mechanisms that may underpin estrogen-mediated neuroprotection. To gain insight into these processes, we performed a protein-protein interaction and functional network analysis in STRING on the various protein changes known to occur after estrogen treatment in SCI.

The STRING analysis included 23 proteins that were upregulated by estrogen administration after SCI (Figure 3; Supplementary Table 1). While nine of the included proteins were disconnected nodes, the remaining ones showed a clear network of protein-protein interactions, largely dominated by two central nodes of proto-oncogene tyrosine-protein kinase (Src) and serine/threonine kinase 1 (Akt1) (Figure 3). This network involved proteins implicated in several important biological processes including anatomical structure development (FDR = 2.94e-05) and morphogenesis (FDR = 5.23e-05), nervous system development (FDR = 0.00010), cell junction organization (FDR = 2.94e-05), cell-matrix adhesion (FDR = 6.07e-05), cellular response to growth factor stimulus (FDR = 2.98e-05), and positive regulation of cell migration (FDR = 5.03e-05). The proteins in this network were also localized to the following cellular components: anchoring junction (FDR = 1.13e-06), cell-cell junction (FDR = 3.74e-05), and cell junction (FDR = 8.34e-05). Further, these proteins were implicated in several KEGG pathways that are known to be neuroprotective. The signalling pathways that were significantly upregulated by estrogen administration after SCI included: PI3K-Akt (FDR = 8.14e-05), Rap1 (FDR = 9.70e-05), HIF-1 (FDR = 0.00014), neurotrophin (FDR = 0.00014), VEGF (FDR = 0.00049), cAMP (FDR = 0.00075), Ras (FDR = 0.00095), ErbB (FDR = 0.00098), and glucagon (FDR = 0.0013). Importantly, the estrogen signalling pathway was also identified as being significantly upregulated (FDR = 0.00017).

**Figure 3.**
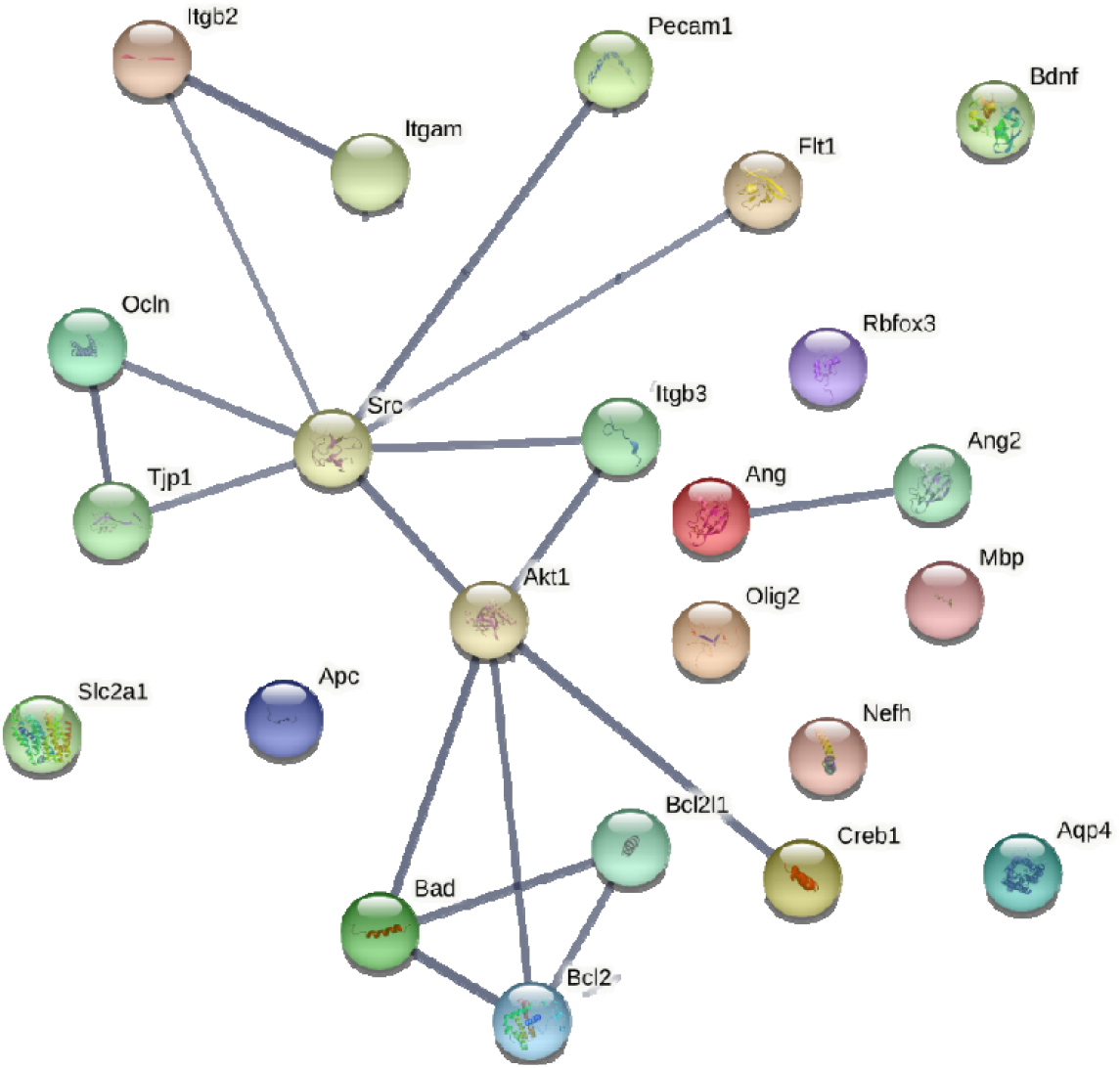
Protein-protein interaction network upregulated by estrogen treatment after SCI. The thickness of the network edges indicates the strength of data support for the interaction.

We also performed a STRING analysis on 28 proteins that were downregulated by estrogen administration after SCI (Figure 4; Supplementary Table 1). Similar to the upregulated proteins, nine of the downregulated proteins were disconnected nodes, with the remainder forming clear protein-protein interaction networks. Unlike the upregulated network, however, the downregulated one was dominated by multiple proteins including ras homolog family member A (RhoA), apoptosis-associated speck-like protein containing a CARD (Pycard, also known as ASC), caspase 1 (Casp1), p38 mitogen-activated protein kinases (MAPKs 11, 12, 13 and 14), MAPK8 (also known as JNK), and Jun (Figure 4). The downregulated proteins in the network involved in myriad positive regulation of biological processes including peptidase activity (FDR = 1.05e-10), cell death (FDR = 1.99e-10), interleukin 1β (IL-1β) production (FDR = 8.10e-10), apoptotic processes (FDR = 8.14e-10), cytokine production (FDR = 3.36e-09), neuron death (FDR = 7.46e-08), and metabolic processes (FDR = 4.60e-07). They were also involved in responses to cytokines (FDR = 2.47e-06), immune system processes (FDR = 6.26e-06), and inflammatory response (FDR = 5.04e-05). Downregulated proteins were localized to three main cellular components including the inflammasome complex (FDR = 2.79e-08), AIM2 inflammasome complex (FDR = 7.93e-05), and the NLRP1 inflammasome complex (FDR = 0.0030). Relative to the upregulated KEGG pathways, there were slightly less downregulated pathways by estrogen administration after SCI, however these tended to be involved in neurodegeneration. The signalling pathways included NOD-like receptor (FDR = 1.92e-20), IL-17 (FDR = 6.92e-14), TNF (FDR = 3.53e-13), Toll like receptor (FDR = 2.58e-10), T cell receptor (FDR= 5.22e-10), apoptosis (FDR = 7.52e-08), P53 (FDR = 1.74e-07), and MAPK (FDR = 3.35e-07).

**Figure 4.**
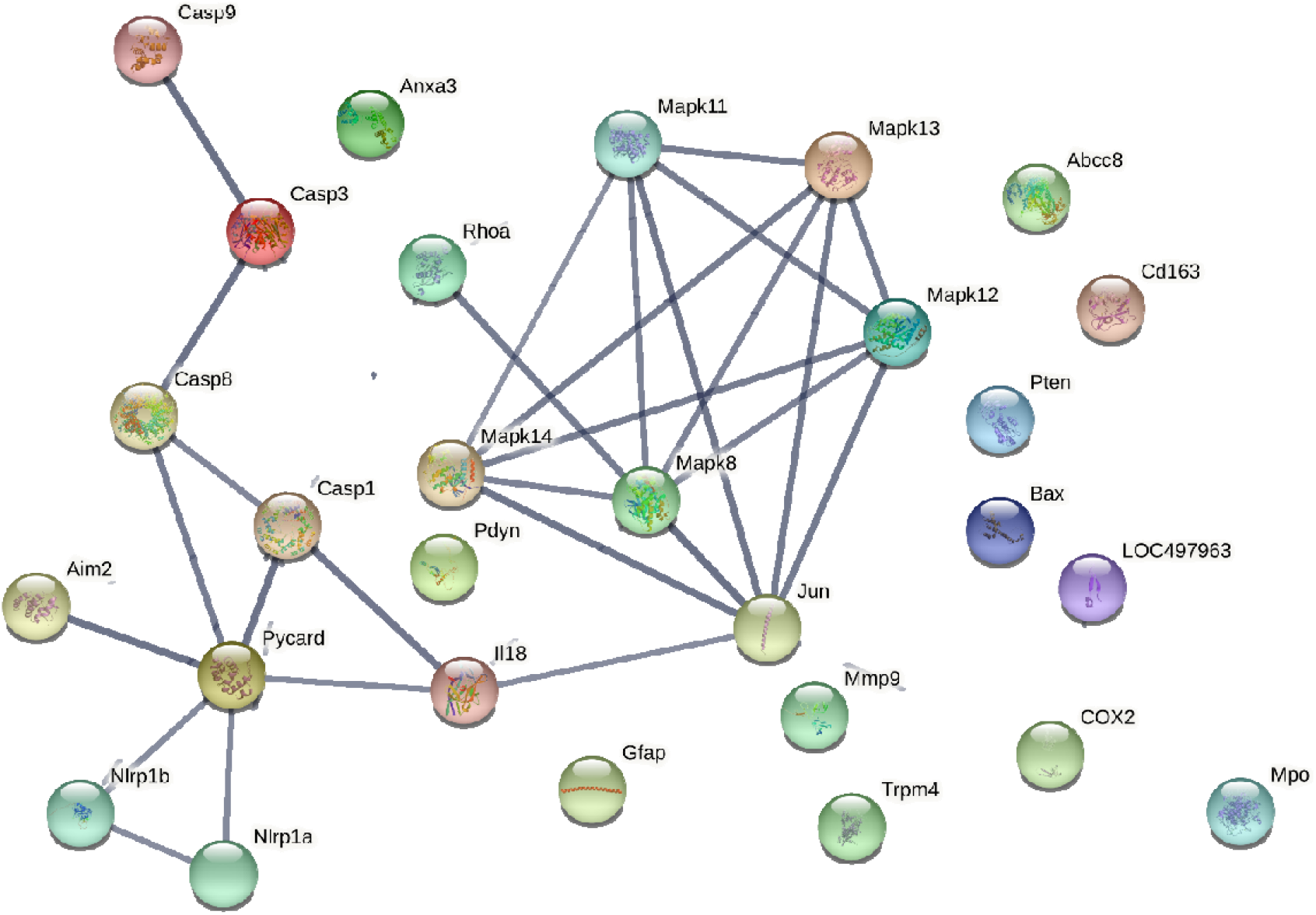
Protein-protein interaction network downregulated by estrogen treatment after SCI. The thickness of the network edges indicates the strength of data support for the interaction.

## 4. Discussion

The present review systematically examined the literature on estrogen and estrogenic compound neuroprotection after SCI. It clearly demonstrated that these types of pharmacological interventions offer protection against SCI. Several estrogens were examined by the included studies: estrogen, estradiol benzoate, Premarin, isopsoralen, genistein, and the SERMs tamoxifen, raloxifene, and bazedoxifene. Broadly, estrogens decreased lesion size, apoptosis, myelin degradation, intracellular calcium influxes and ROS activity after SCI. They also spared white matter and increased cell viability, number of remaining cells, and maintained cell morphology. Combined, this led to clear improvements in functional outcomes after SCI with respect to locomotion and sensation.

### 4.1. Multiple estrogen receptors are involved in neuroprotection and there is likely substantial overlap in their role(s)

Our review also uncovered highly mixed findings with respect to the relative importance of ER subtypes in estrogen and estrogenic compound-mediated neuroprotection. This is not surprising in the context that each of the various outcomes involve different cell types with diverse ontology and likely differing actions mediated by estrogens. Some showed that the ERα and β receptors were critical whereas others implicated the GPER. This finding is in line with previous work. Specifically, ERs are known to have substantial overlap in function and effect and may, to an extent, be able to compensate for the loss of one subtype (Finney et al., 2020; Srivastava et al., 2013). An important line of enquiry, however, that was omitted by the included the studies, is the relative contributions of classical (genomic) and non-classical (non-genomic) ER mechanisms of action to neuroprotection after SCI. For example, estrogen may upregulate VEGF via an ERE upstream from the transcriptional start site to improve angiogenesis after SCI (Mueller et al., 2000; Widenfalk et al., 2003). Further, no studies have examined if a non-classical, non-genomic effect of estrogen is mediated by membrane-bound ERs or not. For example, future studies might be able to use the estrogen-bovine serum albumin conjugate to study the respective roles of membrane and intracellular estrogenic effects after SCI (Taguchi et al., 2004; Temple & Wray, 2005). It is critical that future work continue to establish the specific role of ERs in neuroprotection after SCI and perhaps include cell specific ER modifications.

### 4.2. Estrogen-mediated neuroprotection after SCI likely involves the simultaneous downregulation of pro-inflammatory pathways and upregulation of neuroprotective pathways

Few studies have examined the specific pathways and mechanisms that underly estrogen neuroprotection. The PI3K-Akt pathway has been implicated in estrogen-mediated neuroprotection, with studies showing that administration of the PI3K-Akt inhibitor LY294002 blocks estrogen’s neuroprotective effects after SCI (Chen et al., 2015; Das et al., 2011; Yune et al., 2008). Further, the MAPK pathway has only been partially implicated in estrogenic neuroprotection. One study found that co-administration of estrogen with the MAPK inhibitor PD98059 partially reduced estrogen’s benefits whereas another found that it had no effect (Das et al., 2011; Yune et al., 2008). To date, no high throughput studies of estrogen’s effects on protein changes after SCI have been done. To address this, we performed a STRING functional network analysis on the effect of estrogen on up- and down-regulated proteins after SCI. We found that estrogen recruits several known neuroprotective pathways. This included the PI3K-Akt and MAPK pathway, which is in line with previous estrogen literature as well as a known neuroprotective role for the upregulation of PI3K-Akt and downregulation of MAPK pathway after SCI (Bimbova et al., 2022; Kasuya et al., 2018).

Our STRING analysis also identified several additional pathways that may be critical for estrogen-mediated neuroprotection, most of which have been implicated in SCI more broadly. Estrogen was found to downregulate several pathways that have been shown in SCI to be upregulated after injury including TLR, IL-17, Nod-like signalling, p-53, and T-cell signalling pathways. The TLR pathway is central to the initial induction of the innate immune system after SCI and resultant neuroinflammation (Impellizzeri et al., 2015). The IL-17 and Nod-like signalling pathways play a critical role in SCI-induced inflammation, cytokine release, and apoptosis (Jiang et al., 2017; Zong et al., 2014), and the p53 pathway has been implicated in apoptosis (Saito et al., 2000). Increased T-cell response and signalling after SCI is associated with the development of neuropathic pain as well as the upregulation of cytokines (Costigan et al., 2009). The finding that estrogen downregulates these pathways suggests that estrogen can dampen known neuroinflammatory processes and that this likely plays a role in long-term improved functional and histopathological outcomes after SCI.

In addition to the downregulation of known pro-inflammatory pathways, our STRING analysis showed that estrogen simultaneously upregulates several pathways known to be involved in neuroprotection after SCI. These included Rap1, HIF-1, VEGF, ErbB, and glucagon signalling. Rap1 has been shown to be critical for survival of dorsal root ganglion neurons after treatment with myelin-associated glycoprotein (Taniguchi et al., 2007). Both HIF-1 and VEGF facilitate protection by decreasing inflammation and increasing cell growth (J. Li et al., 2017; Li et al., 2020). ErbB signalling has also been shown to underly Schwann cell-mediated spontaneous remyelination of dorsal column axons after SCI (Bartus et al., 2019). Finally, glucagon signalling in the spinal cord has been implicated in the inhibition of pain hypersensitivity (Gong et al., 2014). These findings suggest that estrogen also can recruit and / or activate established neuroprotective pathways after SCI and highlights the therapeutic potential of estrogen-based therapies.

Interestingly, many of the pathways found to be differentially regulated by estrogen in the present review have been previously implicated in the pathogenesis of SCI using RNA-sequencing (Shi et al., 2017). This suggests that although our STRING analysis was preliminary and used mixed, single results from multiple studies, it likely has significant implications for the specific mechanisms underlying estrogen neuroprotection after SCI. Further, it highlights the delicate nature of estrogen-mediated neuroprotection and the role of simultaneous up- and down-regulation of signalling pathways. As discussed above, however, it is unclear whether estrogen regulates these pathways via genomic or non-genomic mechanisms, and this remains an important consideration for future experimental work to address.

### 4.3. Limitations of the included studies and the present review

An important consideration is that there were many differences in the characteristics of the included studies that may limit the conclusions of the present review. First, studies used a diverse range of SCI models, ranging from contusion to electrolytic, and looked at various timepoints after injury, from immediately after and up to to 42 days later. As such, there may be an underappreciation of the important time-related and injury model-specific pathways and the role of estrogenic therapies within these contexts. Future research would benefit from direct comparisons between injury types and across time of estrogen’s ability to confer neuroprotection after SCI. However, despite the variability in included studies and in cellular diversity we were still able to identify common pathways. This finding may in fact strengthen the conclusion that estrogens are neuroprotective in SCI by modulating diverse pathways and cell types and are likely therapeutically translatable.

The included studies in the present review also used a wide range of estrogen and estrogenic compound doses. Again, although this shows that multiple doses may be effective against the consequences of SCI, the conclusions that we can draw remain limited. Further, no studies performed a human equivalent dose (HED) calculation, indicating an important limitation for these types of studies (Nair & Jacob, 2016). One study, for example, used a dose of tamoxifen that was impossible to achieve in humans, raising questions about the utility and translatability of such findings. This highlights the clear and urgent need for pharmacotherapy studies to consistently use an HED calculation to ensure that the doses being trialled in animals are within the therapeutic range in humans, assuming similar rodent physiology in the examined pathway.

Third, these preclinical studies were limited in their experimental groups. Few studies used female animals and, when they did, they were typically ovariectomized. While ovariectomized females can be a valuable model for studying whether drugs are effective after menopause, it does potentially rule out drug effects in females of childbearing ages. In Australia, for example, around 10% of SCI are in females under the age of 44 (AIHW, 2021). Future studies, therefore, should focus on the use of intact female animals to identify whether estrogen and estrogenic compounds are still neuroprotective against SCI. Most of the included studies also only used animals that were quite young, with only one study comparing treatment effects in young versus old animals (Chaovipoch et al., 2005). Recent research indicated that the incidence of SCI among the elderly is increasing and that they have differing pathophysiological and functional outcomes relative to their younger counterparts with SCI, including a doubled risk of death (Wilson et al., 2020). It is important, therefore, that preclinical models examine age as an important co-variable in determining if estrogen and its compounds confer neuroprotection in SCI.

Finally, the above limitations and variability in experimental parameters prevented us from performing any meta-analyses on the included results in this review. Further, studies often did not include detailed statistical information (e.g. means) that would be required for such analyses. To mitigate any outcome bias associated with reviewing this literature base, we performed a systematic review with clear inclusion and exclusion criteria and reported the methods and results as they were reported in their respective papers. However, there is a need to begin to standardize experimental parameters for preclinical pharmacotherapy studies and for journals to require detailed statistical information in their publications to allow for meta-analyses and establishing effect sizes for estrogenic neuroprotection after SCI.

### 4.4. Considerations for clinical translatability of estrogenic therapies after SCI

Although the present review demonstrated that there is clear preclinical evidence that estrogens are neuroprotective against SCI it is important to note that clinical use of estrogens is not without risk and controversy. More specifically, there is a significant risk profile associated with the prolonged use of estrogens including increased risk of coronary heart disease, thromboembolic events, breast cancer and uterine cancer in women (Lobo, 2017). This suggests that the possible neuroprotective benefits of estrogen treatment after SCI may not outweigh the risks associated with its use. One consideration, however, is that several of the included studies used very short-term or pro re nata (*prn*) administration of estrogen and still found neuroprotective benefits. As such, this type of use may offer the ideal balance between risks and benefits in patients with SCI. Any clinical trials, therefore, that consider the use of estrogens after SCI should focus on *prn* and short-term estrogen therapies.

It is also important to note that our review also showed clear evidence that estrogenic compounds, including SERMs, conferred neuroprotection after SCI. SERMs are a viable alternate strategy to estrogen as they are known to maintain pro-estrogenic benefits while minimizing the risk of side effects and adverse events (Duterte & Smith, 2000). Further, they are already approved for use in humans. In the present review, SERMs studied in the context of SCI included tamoxifen, raloxifene, and bazedoxifene. Generally, SERM neuroprotective benefits after SCI were similar to those reported for estrogen however well-controlled comparative studies are needed to confirm this. It is also important to note that chronic use of tamoxifen has been associated with increased risk of stroke and thromboembolic events in humans (Braithwaite et al., 2003; Bushnell & Goldstein, 2004; Vogel et al., 2006). Newer generation SERMs, including raloxifene and bazedoxifene, have had some success in overcoming this risk profile, such as reducing the risk of thromboembolic events (Vestergaard & Thomsen, 2009; Vogel et al., 2006). SERM use after SCI, therefore, may also be limited to short-term or *prn* administration although it is worth noting that raloxifene and bazedoxifene both showed neuroprotective benefit after SCI.

## 5. Conclusion

SCI is a severe and debilitating condition for which there are few treatment options. One line of pharmacotherapy that may effectively mitigate the effects of SCI are estrogen and estrogenic compounds, like SERMs. Based on the evidence presented in this review, there is substantial pre-clinical evidence that both are neuroprotective and improve the functional outcome of SCI. Although the present review offered some insight into the mechanisms and protein networks that may underlie this neuroprotection, there is a need for future research to provide experimental evidence of these. Further, there is a need for future studies to use consistent SCI models as well as pharmacotherapy doses that fall within the therapeutic range in humans. In doing so, we will begin to have a clear understanding of the role of estrogen, and estrogenic compounds, in neuroprotection after SCI and move toward clinical translatability of these findings.

## Acknowledgements

Figures in the present review were made using Biorender.com and STRING.

