## Supplementary Table for "The neuroprotective effects of estrogen and estrogenic compounds in spinal cord injury"

Supplementary Table 1. Effects of estrogen and estrogenic compounds on RNA expression, protein, and phosphorylation changes after SCI

| **Protein or Gene** | **Role** | **Drug** | **Dose / Concentration** | **Number of Doses** | **Pre or Post SCI** | **Model(s)** | **Effect of Drug** | **References** |
| --- | --- | --- | --- | --- | --- | --- | --- | --- |
| Abcc8 mRNA | Modulator of ATP-sensitive potassium channels and insulin release | Estrogen | 300µg/kg | 3 | Post | Rat, contusion | Temporarily decreased | (Lee et al., 2015) |
| Abcc8* |  | Estrogen | 300µg/kg | 3 | Post | Rat, contusion | Decreased | (Lee et al., 2015) |
| Aim2 mRNA | Control of cell proliferation | Estrogen | 25µg/kg | 6 | Post | Rat, contusion | No effect | (Majidpoor et al., 2020) |
| Aim2* |  | Estrogen | 25µg/kg | 6 | Post | Rat, contusion | Decreased | (Majidpoor et al., 2020) |
| Akt | Cell proliferation, survival, metabolism, angiogenesis | Isopsoralen | 5mg/kg | 1 | Post | Mouse, compression | No effect | (Li et al., 2017) |
|  |  |  | 10mg/kg | 1 | Post | Mouse, compression | No effect | (Li et al., 2017) |
| Akt* (phosphorylated) |  | Estrogen | 1µg | 14 | Post | Mouse, compression | Increased | (Cheng et al., 2016) |
|  |  |  | 100µg/kg | 1 | Post | Rat, contusion | Increased | (Yune et al., 2008) |
|  |  |  | 100nM | Bath | Pre | Motoneuron cells | Increased | (Smith et al., 2009) |
|  |  |  | 150nM | Bath | Post | Motoneuron cells | Increased | (Das et al., 2011) |
|  |  |  | Not specified | Bath | Post | Motoneuron cells | Increased | (Chen et al., 2015) |
|  |  | G1 | 1µg | 14 | Post | Mouse, compression | Increased | (Cheng et al., 2016) |
|  |  |  | 10nM | Bath | Post | Motoneuron cells | Increased | (Chen et al., 2015) |
|  |  | PPT | 50nM | Bath | Post | Motoneuron cells | Increased | (Das et al., 2011) |
|  |  | DPN | 50nM | Bath | Post | Motoneuron cells | Increased | (Das et al., 2011) |
|  |  | Isopsoralen | 5mg/kg | 1 | Post | Mouse, compression | Increased | (Li et al., 2017) |
|  |  |  | 10mg/kg | 1 | Post | Mouse, compression | Increased | (Li et al., 2017) |
|  |  | Premarin | 100µg/kg | 8 | Post | Rat, contusion | Increased | (Haque et al., 2022) |
| Ang-1* | Vascular development and angiogenesis | Estrogen | 10µg/kg | 8 | Post | Rat, contusion | Increased | (Haque et al., 2022) |
|  |  | Premarin | 100µg/kg | 8 | Post | Rat, contusion | Increased | (Haque et al., 2022) |
| Ang-2* | Increase vasopressin | Estrogen | 10µg | 7 | Post | Rat, contusion | Increased | (Samantaray et al., 2016) |
| APC* | Cell migration and adhesion, transcriptional activation, apoptosis | Estrogen | 25µg/kg | 7 | Post | Rat, compression | Increased | (Zendedel et al., 2018) |
|  |  | Tamoxifen | 1mg/day | Implant | Post | Rat, contusion | Increased | (Guptarak et al., 2014) |
| ASC mRNA | Caspase activation | Estrogen | 25µg/kg | 7 | Post | Rat, compression | Decreased | (Zendedel et al., 2018) |
| ASC* |  | Estrogen | 25µg/kg | 6 or 7 | Post | Rat, contusion  Rat, compression | Decreased | (Majidpoor et al., 2020; Majidpoor et al., 2021; Zendedel et al., 2018) |
| Atg5 | Autophagy, mitochondrial quality, lymphocyte development and proliferation, apoptosis | Estradiol benzoate | 0.5mg/kg | 16-17 | Pre and Post | Rat, compression | Decreased | (Lin et al., 2016) |
| Atg7 | Autophagy, axon membrane trafficking, axonal homeostasis, mitophagy | Estradiol benzoate | 0.5mg/kg | 16-17 | Pre and Post | Rat, compression | Decreased | (Lin et al., 2016) |
| AQP4* | Water-selective channel | Tamoxifen | 1mg/day | Implant | Post | Rat, contusion | Increased | (Guptarak et al., 2014) |
| BAD* (phosphorylated) | Positive regulator of cell apoptosis | Estrogen | 150nM | Bath | Post | Motoneuron cells | Increased | (Das et al., 2011) |
|  |  | PPT | 50nM | Bath | Post | Motoneuron cells | Increased | (Das et al., 2011) |
|  |  | DPN | 50nM | Bath | Post | Motoneuron cells | Increased | (Das et al., 2011) |
| Bax* | Apoptosis activator | Estrogen | 1µg | 14 | Post | Mouse, compression | Decreased | (Cheng et al., 2016) |
|  |  |  | 5µg | Implant | Post | Rat, contusion | Decreased | (Haque et al., 2021) |
|  |  |  | 0.008mg/kg | Implant | Post | Rat, contusion | No effect | (Kachadroka et al., 2010) |
|  |  |  | 0.08mg/kg | Implant | Post | Rat, contusion | No effect | (Kachadroka et al., 2010) |
|  |  |  | 300µg/kg | 3 | Pre and Post | Mouse, compression | Decreased | (Cuzzocrea et al., 2008) |
|  |  |  | 0.8mg/kg | Implant | Post | Rat, contusion | Decreased | (Kachadroka et al., 2010) |
|  |  | G1 | 1µg | 14 | Post | Mouse, compression | Decreased | (Cheng et al., 2016) |
|  |  | Isopsoralen | 5mg/kg | 1 | Post | Mouse, compression | Decreased | (Li et al., 2017) |
|  |  |  | 10mg/kg | 1 | Post | Mouse, compression | Decreased | (Li et al., 2017) |
|  |  | Premarin | 100µg/kg | 8 | Post | Rat, contusion | Decreased | (Haque et al., 2022) |
| Bcl-2* | Blocks apoptotic death | Estrogen | 1µg | 14 | Post | Mouse, compression | Increased | (Cheng et al., 2016) |
|  |  |  | 3µg/kg | 1 | Pre | Rat, contusion | Increased | (Yune et al., 2004) |
|  |  |  | 5µg | Implant | Post | Rat, contusion | Increased | (Haque et al., 2021) |
|  |  |  | 100µg/kg | 1 | Post | Rat, contusion | Increased | (Yune et al., 2004; Yune et al., 2008) |
|  |  |  | 300µg/kg | 1 or 3 | Pre, Pre and Post | Rat, contusion  Mouse, compression | Increased | (Cuzzocrea et al., 2008; Yune et al., 2004) |
|  |  |  | 150nM | Bath | Post | Motoneuron cells | Increased | (Das et al., 2011) |
|  |  | G1 | 1µg | 14 | Post | Mouse, compression | Increased | (Cheng et al., 2016) |
|  |  | PPT | 50nM | Bath | Post | Motoneuron cells | Increased | (Das et al., 2011) |
|  |  | DPN | 50nM | Bath | Post | Motoneuron cells | Increased | (Das et al., 2011) |
|  |  | Isopsoralen | 5mg/kg | 1 | Post | Mouse, compression | Increased | (Li et al., 2017) |
|  |  |  | 10mg/kg | 1 | Post | Mouse, compression | Increased | (Li et al., 2017) |
|  |  | Premarin | 100µg/kg | 8 | Post | Rat, contusion | Increased | (Haque et al., 2022) |
| Bcl-Xl/Bcl-Xs | Apoptotic regulator | Estrogen | 0.008mg/kg | Implant | Post | Rat, contusion | No effect | (Kachadroka et al., 2010) |
|  |  |  | 0.08mg/kg | Implant | Post | Rat, contusion | No effect | (Kachadroka et al., 2010) |
|  |  |  | 100µg/kg | 1 | Pre | Rat, contusion | Increased | (Yune et al., 2004) |
|  |  |  | 0.8mg/kg | Implant | Post | Rat, contusion | Increased | (Kachadroka et al., 2010) |
| BDNF* | Neuronal survival and growth | Estrogen | 1µg | 14 | Post | Mouse, compression | Increased | (Cheng et al., 2016) |
|  |  | G1 | 1µg | 14 | Post | Mouse, compression | Increased | (Cheng et al., 2016) |
| Beclin1 mRNA | Regulates autophagy | Estradiol benzoate | 0.5mg/kg | 16-17 | Pre and Post | Rat, compression | Decreased | (Lin et al., 2016) |
| Beclin1 |  | Estradiol benzoate | 0.5mg/kg | 16-17 | Pre and Post | Rat, compression | Decreased | (Lin et al., 2016) |
| Calpain | Intracellular cysteine protease | Estrogen | 4mg/kg | 2 | Post | Rat, contusion | No effect | (Sribnick et al., 2006) |
| Capase-1 mRNA | Apoptosis activator; IL-1 activator | Estrogen | 25µg/kg | 6 | Post | Rat, contusion | Decreased | (Majidpoor et al., 2020) |
| Caspase-1* |  | Estrogen | 25µg/kg | 6 | Post | Rat, contusion | Decreased | (Majidpoor et al., 2020) 31 |
| Caspase-3* | Execution-phase of cell apoptosis | Estrogen | 1µg | 14 | Post | Mouse, compression | Decreased | (Cheng et al., 2016) |
|  |  |  | 25µg/kg | 7 | Post | Rat, compression | Decreased | (Zendedel et al., 2018) |
|  |  |  | 100µg/kg | 1 | Pre | Rat, contusion | Decreased | (Yune et al., 2004) |
|  |  |  | 300µg/kg | 3 | Post | Rat, contusion | Decreased | (Lee et al., 2012) |
|  |  | G1 | 1µg | 14 | Post | Mouse, compression | Decreased | (Cheng et al., 2016) |
|  |  | Premarin | 1mg/kg | Implant | Post | Rat, compression | Decreased | (Chen et al., 2010) |
|  |  | Genistein | 50nM | Bath | Post | Motoneuron cells | Decreased | (McDowell et al., 2011) |
|  |  | Bazedoxifene | 1mg/kg | 2 | Pre and Post | Rat, compression | Decreased | (Kim et al., 2021) |
| Caspase-8* | Programmed cell death | Estrogen | 150nM | Bath | Post | Motoneuron cells | Decreased | (Das et al., 2011) |
|  |  | PPT | 50nM | Bath | Post | Motoneuron cells | Decreased | (Das et al., 2011) |
|  |  | DPN | 50nM | Bath | Post | Motoneuron cells | Decreased | (Das et al., 2011) |
|  |  | Genistein | 50nM | Bath | Post | Motoneuron cells | Decreased | (McDowell et al., 2011) |
| Caspase 9* | Apoptosis activator | Estrogen | 300µg/kg | 3 | Post | Rat, contusion | Decreased | (Lee et al., 2012) |
|  |  |  | 150nM | Bath | Post | Motoneuron cells | Decreased | (Das et al., 2011) |
|  |  | PPT | 50nM | Bath | Post | Motoneuron cells | Decreased | (Das et al., 2011) |
|  |  | DPN | 50nM | Bath | Post | Motoneuron cells | Decreased | (Das et al., 2011) |
|  |  | Genistein | 50nM | Bath | Post | Motoneuron cells | Decreased | (McDowell et al., 2011) |
| Cd11 | Adhesion between cells | Estrogen | 0.5mg | Implant | Pre, Post | Mouse, transection  Rat, contusion | No effect | (Siriphorn et al., 2012; Webb et al., 2006) |
|  |  |  | 5mg | Implant | Post | Rat, contusion | Decreased | (Siriphorn et al., 2012) |
| Cd11b* | α chain of integrin receptor | Estrogen | 1µg/kg | Implant | Post | Rat, contusion | Decreased | (Samantaray et al., 2011) |
|  |  |  | 5µg/kg | Implant | Post | Rat, contusion | Decreased | (Samantaray et al., 2011) |
|  |  |  | 10µg | 7 | Post | Rat, contusion | Decreased | (Samantaray et al., 2016) |
|  |  |  | 10µg/kg | Implant | Post | Rat, contusion | Decreased | (Samantaray et al., 2011) |
|  |  |  | 4mg/kg | 2 or 7 | Post | Rat, contusion | Decreased | (Sribnick et al., 2010; Sribnick et al., 2005) |
|  |  | Tamoxifen | 5mg/kg | 1 | Post | Rat, contusion | Decreased | (Tian et al., 2009) |
| Cd13 | Transmembrane metalloprotease | Premarin | 100µg/kg | 8 | Post | Rat, contusion | Increased | (Haque et al., 2022) |
| Cd31* | Cell adhesion | Estrogen | 10µg | 7 | Post | Rat, contusion | Increased | (Samantaray et al., 2016) |
|  |  |  | 25µg/kg | 7 | Post | Rat, contusion | Increased | (Tan et al., 2020) |
| Cd61* | Integrin binding and cell adhesion | Estrogen | 25µg/kg | 7 | Post | Rat, contusion | Increased | (Tan et al., 2020) |
| Cd68 | Phagocytosis of tissue macrophages | Estrogen | 2ng/day | Implant | Post | Mouse, contusion | No effect | (Gottipati et al., 2021) |
|  |  |  | 0.1mg/kg | 1 | Post | Rat, compression | No effect | 40 |
|  |  |  | 300µg/kg | 3 | Post | Rat, contusion | Decreased | (Lee et al., 2015) |
|  |  |  | 4mg/kg | 1 | Post | Rat, compression | Temporarily decreased | (Ritz & Hausmann, 2008) |
|  |  | Tamoxifen | 300mg/kg | 4 | Post | Mouse, contusion | No effect | (Pukos & McTigue, 2020) |
| Cd163* | Clearance and endocytosis of hemoglobin/haptoglobin complexes by macrophages | Estrogen | 4mg/kg | 2 or 7 | Post | Rat, contusion | Decreased | (Sribnick et al., 2010; Sribnick et al., 2005) |
| CGRP | Vasodilation | Estrogen | 0.5mg | Implant | Pre | Mouse, transection | No effect | (Webb et al., 2006) |
| c-Jun* (phosphorylated) | Positive regulation of transcription | Estrogen | 300µg/kg | 1 or 3 | Post | Rat, contusion | Decreased | (Lee et al., 2018; Lee et al., 2012) |
| CNPase | Hydrolysis of cyclic nucleotides to monophosphates | Bazedoxifene | 1mg/kg | 8 | Pre and Post | Rat, compression | No effect | (Kim et al., 2021) |
|  |  | Tamoxifen | 1mg/day | Implant | Post | Rat, contusion | Increased | (Guptarak et al., 2014) |
| COX-2 mRNA | Mitochondrial electron transport; positive regulation of vasoconstriction | Estrogen | 300µg/kg | 3 | Pre and Post, Post | Mouse, compression  Rat, contusion | Decreased  Temporarily decreased | (Cuzzocrea et al., 2008; Lee et al., 2015) |
| Cox-2* |  | Estrogen | 300µg/kg | 1 or 3 | Post | Rat, contusion | Decreased | (Lee et al., 2018; Lee et al., 2015) |
|  |  |  | 4mg/kg | 7 | Post | Rat, contusion | Decreased | (Sribnick et al., 2010) |
|  |  | Genistein | 50nM | Bath | Post | Motoneuron cells | Decreased | (McDowell et al., 2011) |
|  |  | Premarin | 100µg/kg | 8 | Post | Rat, contusion | Decreased | (Haque et al., 2022) |
| CRE | DNA recombination | Estrogen | 100µg/kg | 1 | Post | Rat, contusion | Increased | (Yune et al., 2008) |
| CREB* (phosphorylated) | cAMP pathway transcription factor | Estrogen | 100µg/kg | 1 | Post | Rat, contusion | Increased | (Yune et al., 2008) |
|  |  |  | 150nM | Bath | Post | Motoneuron cells | Increased | (Das et al., 2011) |
|  |  | PPT | 50nM | Bath | Post | Motoneuron cells | Increased | (Das et al., 2011) |
|  |  | DPN | 50nM | Bath | Post | Motoneuron cells | Increased | (Das et al., 2011) |
| Cytochrome c | Mitochondrial electron transport chain; initiation of apoptosis | Estrogen | 4mg/kg | 2 | Post | Rat, contusion | No effect | (Sribnick et al., 2006) |
|  |  |  | 150nM | Bath | Post | Motoneuron cells | Increased | (Das et al., 2011) |
|  |  | PPT | 50nM | Bath | Post | Motoneuron cells | Increased | (Das et al., 2011) |
|  |  | DPN | 50nM | Bath | Post | Motoneuron cells | Increased | (Das et al., 2011) |
|  |  | Genistein | 50nM | Bath | Post | Motoneuron cells | Decreased | (McDowell et al., 2011) |
| ERK (phosphorylated) | Proliferation, differentiation, transcription regulation, development | Estrogen | 1µg | 14 | Post | Mouse, compression | Increased | (Cheng et al., 2016) |
|  |  |  | 100µg/kg | 1 | Post | Rat, contusion | Increased | (Yune et al., 2008) |
|  |  |  | 300µg/kg | 1 | Post | Rat, contusion | Decreased | (Lee et al., 2018) |
|  |  |  | 150nM | Bath | Post | Motoneuron cells | Decreased | (Das et al., 2011) |
|  |  | G1 | 1µg | 14 | Post | Mouse, compression | Increased | (Cheng et al., 2016) |
|  |  | PPT | 50nM | Bath | Post | Motoneuron cells | Decreased | (Das et al., 2011) |
|  |  | DPN | 50nM | Bath | Post | Motoneuron cells | Decreased | (Das et al., 2011) |
| Flk-1 | Receptor for VEGF | Estrogen | 10µg/kg | 8 | Post | Rat, contusion | No effect | (Haque et al., 2022) |
|  |  | Premarin | 100µg/kg | 8 | Post | Rat, contusion | Increased | (Haque et al., 2022) |
| Flt-1* | Receptor for VEGF | Estrogen | 10µg/kg | 8 | Post | Rat, contusion | Increased | (Haque et al., 2022) |
|  |  | Premarin | 100µg/kg | 8 | Post | Rat, contusion | Increased | (Haque et al., 2022) |
| Gap43 | Development and axon regeneration | Estrogen | 25µg/kg | 7 | Post | Rat, contusion | Temporarily increased | (Tan et al., 2020) |
| GDNF | Transcription factor | Premarin | 1mg/kg | Implant | Post | Rat, compression | Increased | (Chen et al., 2010) |
| GFAP* | Intermediate filament protein of mature astrocytes | Estrogen | 5µg | Implant | Post | Rat, contusion | Decreased | (Haque et al., 2021) |
|  |  |  | 10µg | 7 | Post | Rat, contusion | Decreased | (Samantaray et al., 2016) |
|  |  |  | 100µg | 7 | Post | Rat, contusion | Decreased | (Samantaray et al., 2016) |
|  |  |  | 0.1mg/kg | 1 | Post | Rat, compression | Temporarily increased | (Ritz & Hausmann, 2008) |
|  |  |  | 300µg/kg | 1 | Post | Rat, contusion | Decreased | (Lee et al., 2018) |
|  |  |  | 0.5mg | Implant | Post | Rat, contusion | No effect | (Siriphorn et al., 2012) |
|  |  |  | 4mg/kg | 1 or 7 | Post | Rat, compression  Rat, contusion | Temporarily increased,  Decreased | (Ritz & Hausmann, 2008; Sribnick et al., 2010) |
|  |  |  | 5mg | Implant | Post | Rat, contusion | Decreased | (Siriphorn et al., 2012) |
|  |  | Premarin | 100µg/kg | 8 | Post | Rat, contusion | Decreased | (Haque et al., 2022) |
|  |  |  | 1mg/kg | Implant | Post | Rat, compression | Decreased | (Chen et al., 2010) |
|  |  | Tamoxifen | 0.71mg/day | Implant | Post | Rat, contusion | Decreased | (Colón et al., 2018; Colón et al., 2016) |
|  |  |  | 1mg/day | Implant | Post | Rat, contusion | Decreased | (Guptarak et al., 2014) |
|  |  |  | 300mg/kg | 4 | Post | Mouse, contusion | No effect | (Pukos & McTigue, 2020) |
| Glt-1 | Glutamate transporter protein | Estrogen | 0.08mg/kg | Implant | Post | Rat, contusion | Temporarily increased | (Olsen et al., 2010) |
| Glut1* | Glucose transporter | Estrogen | 10µg | 7 | Post | Rat, contusion | Increased | (Samantaray et al., 2016) |
|  |  | Premarin | 100µg/kg | 8 | Post | Rat, contusion | Increased | (Haque et al., 2022) |
| GROα mRNA | Growth factor | Estrogen | 300µg/kg | 3 | Post | Rat, contusion | Temporarily decreased | (Lee et al., 2015) |
| GSTπ | Detoxification | Tamoxifen | 300mg/kg | 4 | Post | Mouse, contusion | No effect | (Pukos & McTigue, 2020) |
| Iba1 | Binds actin and calcium; macrophage activation | Estrogen | 5µg | Implant | Post | Rat, contusion | Decreased | (Haque et al., 2021) |
|  |  |  | 25µg/kg | 7 | Post | Rat, compression | Increased | (Zendedel et al., 2018) |
| IκBα | NF-κB inflammatory response | Estrogen | 4mg/kg | 2 | Post | Rat, contusion | No effect | (Sribnick et al., 2005) |
| IL-1α | Pleiotropic cytokine; immune response; inflammation | Estrogen | 0.1mg/kg | 1 | Post | Rat, compression | No effect | (Ritz & Hausmann, 2008) |
|  |  |  | 4mg/kg | 1 | Post | Rat, compression | Temporarily increased | (Ritz & Hausmann, 2008) |
| IL-1β mRNA | Cytokine; inflammatory response; cell proliferation, differentiation, and apoptosis | Estrogen | 25µg/kg | 7 | Post | Rat, compression | Decreased | (Zendedel et al., 2018) |
|  |  |  | 300µg/kg | 3 | Pre and Post | Mouse, compression | Decreased | (Cuzzocrea et al., 2008) |
| IL-1β |  | Estrogen | 25µg/kg | 6 or 7 | Post | Rat, contusion  Rat, compression | Decreased | (Majidpoor et al., 2020; Majidpoor et al., 2021; Zendedel et al., 2018) |
|  |  |  | 0.1mg/kg | 1 | Post | Rat, compression | No effect | (Ritz & Hausmann, 2008) |
|  |  |  | 300µg/kg | 1 or 3 | Post | Rat, contusion | Decreased  Temporarily decreased | (Lee et al., 2018; Lee et al., 2015) |
|  |  |  | 4mg/kg | 1 | Post | Rat, compression | Temporarily increased | (Ritz & Hausmann, 2008) |
|  |  | Raloxifene | 30mg/kg | 1 | Post | Rat, compression | Decreased | (Ismailoglu et al., 2013) |
|  |  | Tamoxifen | 5mg/kg | 1 | Post | Rat, compression  Rat, contusion | Decreased | (Ismailoglu et al., 2010; Tian et al., 2009) |
| IL-6 mRNA | Cytokine; inflammation; maturation of B cells | Estrogen | 300µg/kg | 3 | Pre and Post | Mouse, compression | Decreased | (Cuzzocrea et al., 2008) |
| IL-6 |  | Estrogen | 0.1mg/kg | 1 | Post | Rat, compression | No effect | (Ritz & Hausmann, 2008) |
|  |  |  | 300µg/kg | 1 | Post | Rat, contusion | Decreased | (Lee et al., 2018) |
|  |  |  | 4mg/kg | 1 | Post | Rat, compression | No effect | (Ritz & Hausmann, 2008) |
|  |  | Bazedoxifene | 1mg/kg | 3 | Pre and Post | Rat, compression | Decreased | (Kim et al., 2021) |
| IL-18 mRNA | Proinflammatory cytokine | Estrogen | 25µg/kg | 7 | Post | Rat, compression | Decreased | (Zendedel et al., 2018) |
| IL-18* |  | Estrogen | 25µg/kg | 6 or 7 | Post | Rat, contusion  Rat, compression | Decreased | (Majidpoor et al., 2020; Zendedel et al., 2018) |
| iNOS mRNA | Reactive free radical | Estrogen | 300µg/kg | 3 | Pre and Post, Post | Mouse, compression  Rat, contusion | Decreased | (Cuzzocrea et al., 2008; Lee et al., 2015) |
| iNos* |  | Estrogen | 300µg/kg | 1 or 3 | Post | Rat, contusion | Decreased | (Lee et al., 2018; Lee et al., 2015) |
| JNK1/2 | Mediates immediate early gene response | Estrogen | 300µg/kg | 3 | Post | Rat, contusion | No effect | (Lee et al., 2012) |
| JNK* (phosphorylated) |  | Estrogen | 300µg/kg | 1 | Post | Rat, contusion | Decreased | (Lee et al., 2018) |
|  |  |  | 4mg/kg | 1 | Post | Rat, contusion | Decreased | (Rong et al., 2012) |
| Kir4.1 | Inward rectifier-type potassium channel | Estrogen | 0.08mg/kg | Implant | Post | Rat, contusion | Temporarily increased | (Olsen et al., 2010) |
| Lc3 mRNA | Light chain subunit of either MAP1A or MAP1B | Estradiol benzoate | 0.5mg/kg | 16-17 | Pre and Post | Rat, compression | Decreased | (Lin et al., 2016) |
| Lc3* |  | Estradiol benzoate | 0.5mg/kg | 16-17 | Pre and Post | Rat, compression | Decreased | (Lin et al., 2016) |
| MAG mRNA | Myelination | Tamoxifen | 5mg/kg | 1 | Post | Rat, contusion | Decreased | (Tian et al., 2009) |
| p38 MAPK* (phosphorylated) | Cell differentiation, apoptosis, and autophagy | Estrogen | 300µg/kg | 1 | Post | Rat, contusion | Decreased | (Lee et al., 2018) |
| MBP* | Constituent of myelin sheath of oligodendrocytes and Schwann cells | Estrogen | 5µg | Implant | Post | Rat, contusion | Increased | (Haque et al., 2021) |
|  |  |  | 4mg/kg | 1 | Post | Rat, electrolytic | Increased | (Afhami et al., 2016) |
|  |  | Premarin | 100µg/kg | 8 | Post | Rat, contusion | Increased | (Haque et al., 2022) |
|  |  | Tamoxifen | 1mg/day | Implant | Post | Rat, contusion | Increased | (Guptarak et al., 2014) |
| MCP-1 mRNA | Cytokine; immunoregulatory and inflammatory processes | Estrogen | 300µg/kg | 3 | Pre and Post, Post | Mouse, compression  Rat, contusion | Decreased  Temporary decreased | (Cuzzocrea et al., 2008; Lee et al., 2015) |
| Mip-1α mRNA | Inflammatory responses | Estrogen | 300µg/kg | 3 | Post | Rat, contusion | No effect | (Lee et al., 2015) |
| Mip-1β mRNA | Mitogen-inducible monokine; inflammatory responses | Estrogen | 300µg/kg | 3 | Post | Rat, contusion | Temporarily decreased | (Lee et al., 2015) |
| Mip-2α mRNA | Immunoregulatory and inflammatory processes | Estrogen | 300µg/kg | 3 | Post | Rat, contusion | Temporarily decreased | (Lee et al., 2015) |
| miR106a | Post-transcriptional regulation of gene expression | Estrogen | 150nM | Bath | Post | Motoneuron cells | No effect | (Chakrabarti et al., 2014) |
|  |  | PPT | 50nM | Bath | Post | Motoneuron cells | No effect | (Chakrabarti et al., 2014) |
|  |  | WAY200070 | 100nM | Bath | Post | Motoneuron cells | No effect | (Chakrabarti et al., 2014) |
| miR-17 | Post-transcriptional regulation of gene expression | Estrogen | 150nM | Bath | Post | Motoneuron cells | Increased | (Chakrabarti et al., 2014) |
|  |  | PPT | 50nM | Bath | Post | Motoneuron cells | Increased | (Chakrabarti et al., 2014) |
|  |  | WAY200070 | 100nM | Bath | Post | Motoneuron cells | Increased | (Chakrabarti et al., 2014) |
| miR-206 | Post-transcriptional regulation of gene expression | Estrogen | 150nM | Bath | Post | Motoneuron cells | Increased | (Chakrabarti et al., 2014) |
|  |  | PPT | 50nM | Bath | Post | Motoneuron cells | No effect | (Chakrabarti et al., 2014) |
|  |  | WAY200070 | 100nM | Bath | Post | Motoneuron cells | Increased | (Chakrabarti et al., 2014) |
| miR-21 | Post-transcriptional regulation of gene expression | Estrogen | 150nM | Bath | Post | Motoneuron cells | No effect | (Chakrabarti et al., 2014) |
|  |  | PPT | 50nM | Bath | Post | Motoneuron cells | No effect | (Chakrabarti et al., 2014) |
|  |  | WAY200070 | 100nM | Bath | Post | Motoneuron cells | No effect | (Chakrabarti et al., 2014) |
| miR-7-1 | Post-transcriptional regulation of gene expression | Estrogen | 150nM | Bath | Post | Motoneuron cells | Increased | (Chakrabarti et al., 2014) |
|  |  | PPT | 50nM | Bath | Post | Motoneuron cells | Increased | (Chakrabarti et al., 2014) |
|  |  | WAY200070 | 100nM | Bath | Post | Motoneuron cells | Increased | (Chakrabarti et al., 2014) |
| Mmp1 mRNA | Breakdown of extracellular matrix | Estrogen | 300µg/kg | 3 | Post | Rat, contusion | Decreased | (Na et al., 2015) |
| Mmp2 mRNA | Breakdown of extracellular matrix | Estrogen | 300µg/kg | 3 | Post | Rat, contusion | Decreased  No effect | (Lee et al., 2015; Na et al., 2015) |
| Mmp3 mRNA | Breakdown of extracellular matrix | Estrogen | 300µg/kg | 3 | Post | Rat, contusion | Decreased | (Na et al., 2015) |
| Mmp9 mRNA | Breakdown of extracellular matrix | Estrogen | 300µg/kg | 3 | Post | Rat, contusion | Decreased | (Lee et al., 2015; Na et al., 2015) |
| Mmp9* |  | Estrogen | 300µg/kg | 3 | Post | Rat, contusion | Decreased | (Lee et al., 2015) |
| Mmp10 mRNA | Breakdown of extracellular matrix | Estrogen | 300µg/kg | 3 | Post | Rat, contusion | Decreased | (Na et al., 2015) |
| Mmp13 mRNA | Breakdown of extracellular matrix | Estrogen | 300µg/kg | 3 | Post | Rat, contusion | Decreased | (Na et al., 2015) |
| MOG | Immune-mediated demyelination | Bazedoxifene | 1mg/kg | 8 | Pre and Post | Rat, compression | Increased | (Kim et al., 2021) |
|  |  | Tamoxifen | 1mg/day | Implant | Post | Rat, contusion | Increased | (Guptarak et al., 2014) |
| MPO* | Microbial activity of neutrophils | Estrogen | 300µg/kg | 3 | Post | Rat, contusion | Decreased | (Lee et al., 2015) |
| NeuN* | Post-mitotic neuron marker | Estrogen | 2ng/day | Implant | Post | Mouse, contusion | Increased | (Gottipati et al., 2021) |
|  |  | Premarin | 100µg/kg | 8 | Post | Rat, contusion | Increased | (Haque et al., 2022) |
|  |  | Tamoxifen | 0.71mg/day | Implant | Post | Rat, contusion | Increased | (Colón et al., 2018; Colón et al., 2016) |
| NF-H* | Axoskeleton and maintain neuronal calibre | Estrogen | 2ng/day | Implant | Post | Mouse, contusion | Increased | (Gottipati et al., 2021) |
|  |  | Tamoxifen | 0.71mg/day | Implant | Post | Rat, contusion | Increased | (Colón et al., 2016) |
| NF-κB mRNA | Transcription regulation | Estrogen | 25µg/kg | 6 | Post | Rat, contusion | Decreased | (Majidpoor et al., 2020) |
| NF-κB |  | Estrogen | 4mg/kg | 2 | Post | Rat, contusion | No effect | (Sribnick et al., 2005) |
|  |  | Genistein | 50nM | Bath | Post | Motoneuron cells | Decreased | (McDowell et al., 2011) |
|  |  | Premarin | 100µg/kg | 8 | Post | Rat, contusion | Decreased | (Haque et al., 2022) |
| NFP | Nodule organogenesis | Premarin | 100µg/kg | 8 | Post | Rat, contusion | Increased | (Haque et al., 2022) |
| NG2 | Proteoglycan; proliferation and migration of endothelial cells | Bazedoxifene | 1mg/kg | 8 | Pre and Post | Rat, compression | No effect | (Kim et al., 2021) |
|  |  | Tamoxifen | 300mg/kg | 4 | Post | Mouse, contusion | No effect | (Pukos & McTigue, 2020) |
| Nitrotyrosine mRNA | Reactive oxygen species | Estrogen | 300µg/kg | 3 | Pre and Post | Mouse, compression | Decreased | (Cuzzocrea et al., 2008) |
| Nlrc4 mRNA | Innate immune response | Estrogen | 25µg/kg | 7 | Post | Rat, compression | No effect | (Zendedel et al., 2018) |
| Nlrc4 |  | Estrogen | 25µg/kg | 7 | Post | Rat, compression | No effect | (Zendedel et al., 2018) |
| Nlrp1 mRNA | Sensor component of Nlrp1 inflammasome | Estrogen | 25µg/kg | 6 | Post | Rat, contusion | Decreased | (Majidpoor et al., 2020) |
| Nlrp1* |  | Estrogen | 25µg/kg | 6 | Post | Rat, contusion | Decreased | (Majidpoor et al., 2020) |
| Nlrp1b mRNA | Sensor component of Nlrp1b inflammasome | Estrogen | 25µg/kg | 7 | Post | Rat, compression | Decreased | (Zendedel et al., 2018) |
| Nlrp1b * |  | Estrogen | 25µg/kg | 7 | Post | Rat, compression | Decreased | (Zendedel et al., 2018) |
| Nlrp3 mRNA | Sensor component of the Nlrp3 inflammasome | Estrogen | 25µg/kg | 6 or 7 | Post | Rat, contusion  Rat, compression | No effect  Decreased | (Majidpoor et al., 2020; Zendedel et al., 2018) |
| Nlrp3 |  | Estrogen | 25µg/kg | 6 or 7 | Post | Rat, contusion  Rat, compression | No effect,  Decreased | (Majidpoor et al., 2020; Majidpoor et al., 2021; Zendedel et al., 2018) |
| Nogo-A mRNA | Stabilization of endoplasmic reticulum; membrane morphogenesis | Tamoxifen | 5mg/kg | 1 | Post | Rat, contusion | Temporarily decreased | (Tian et al., 2009) |
| Nox2 mRNA | Inflammation regulation | Estrogen | 25µg/kg | 6 | Post | Rat, contusion | Decreased | (Majidpoor et al., 2020) |
| O4 | Transcription factor | Bazedoxifene | 1mg/kg | 8 | Pre and Post | Rat, compression | Increased | (Kim et al., 2021) |
| Occludin* | Formation and regulation of tight junction | Estrogen | 300µg/kg | 3 | Post | Rat, contusion | Increased | (Lee et al., 2015) |
| Olig-2* | Oligodendrocyte and motor neuron specification | Estrogen | 25µg/kg | 7 | Post | Rat, compression | Increased | (Zendedel et al., 2018) |
|  |  | Bazedoxifene | 1mg/kg | 8 | Pre and Post | Rat, compression | Decreased | (Kim et al., 2021) |
|  |  | Tamoxifen | 0.71mg/day | Implant | Post | Rat, contusion | Decreased | (Colón et al., 2018) |
| OMgp | Myelination; cell adhesion | Tamoxifen | 5mg/kg | 1 | Post | Rat, contusion | Decreased | (Tian et al., 2009) |
| κ opioid receptor (phosphor-serine-369) | Receptor for endogenous α-neoendorphins and dynorphins | Estrogen | 2ng/day | Implant | Post | Mouse, contusion | No effect | (Gottipati et al., 2021) |
| PI3K | Regulation of cell survival, migration, differentiation, transcription, and translation | Isopsoralen | 5mg/kg | 1 | Post | Mouse, compression | No effect | (Li et al., 2017) |
|  |  |  | 10mg/kg | 1 | Post | Mouse, compression | No effect | (Li et al., 2017) |
| PI3K (phosphorylated) |  | Isopsoralen | 5mg/kg | 1 | Post | Mouse, compression | Increased | (Li et al., 2017) |
|  |  |  | 10mg/kg | 1 | Post | Mouse, compression | Increased | (Li et al., 2017) |
| Prodynorphin* | Preproprotein precursor of secreted opioid peptides | Estrogen | 0.25mg | Implant | Pre | Rat, contusion | Decreased | (Gupta & Hubscher, 2012) |
| PTEN* | Protein phosphatase | Estrogen | 100nM | Bath | Pre | Motoneuron cells | Decreased | (Smith et al., 2009) |
| RhoA* | GTPase | Estrogen | 300µg/kg | 3 | Post | Rat, contusion | Decreased | (Lee et al., 2012) |
| Src (phosphorylated) | Non-receptor protein tyrosine kinase; gene transcription, immune response, cell adhesion, apoptosis, migration, and transformation | Estrogen | 150nM | Bath | Post | Motoneuron cells | Increased | (Das et al., 2011) |
|  |  | PPT | 50nM | Bath | Post | Motoneuron cells | Increased | (Das et al., 2011) |
|  |  | DPN | 50nM | Bath | Post | Motoneuron cells | Increased | (Das et al., 2011) |
| Sqstm1 | Autophagy receptor | Estradiol benzoate | 0.5mg/kg | 16-17 | Pre and Post | Rat, compression | Decreased | (Lin et al., 2016) |
| tBid (mitochondrial) | Mitochondrial translocation, release of cytochrome c | Estrogen | 150nM | Bath | Post | Motoneuron cells | Decreased | (Das et al., 2011) |
|  |  | PPT | 50nM | Bath | Post | Motoneuron cells | Decreased | (Das et al., 2011) |
|  |  | DPN | 50nM | Bath | Post | Motoneuron cells | Decreased | (Das et al., 2011) |
| Tie-2 | Vascular endothelial function | Premarin | 100µg/kg | 8 | Post | Rat, contusion | Increased | (Haque et al., 2022) |
| Tlr4 mRNA | Inflammatory response | Estrogen | 25µg/kg | 6 | Post | Rat, contusion | Decreased | (Majidpoor et al., 2020) |
| TNF-α mRNA | Cytokine | Estrogen | 300µg/kg | 3 | Pre and Post, Post | Mouse, compression  Rat, contusion | Decreased  Temporarily decreased | (Cuzzocrea et al., 2008; Lee et al., 2015) |
| TNF-α |  | Raloxifene | 30mg/kg | 1 | Post | Rat, compression | Decreased | (Ismailoglu et al., 2013) |
|  |  | Tamoxifen | 5mg/kg | 1 | Post | Rat, compression | Decreased | (Ismailoglu et al., 2010) |
| Trpm4 mRNA | Calcium-activated non-selective cation channel | Estrogen | 300µg/kg | 3 | Post | Rat, contusion | Temporarily decreased | (Lee et al., 2015) |
| Trpm4* |  | Estrogen | 300µg/kg | 3 | Post | Rat, contusion | Decreased | (Lee et al., 2015) |
| Txnip mRNA | Oxidative stress mediator | Estrogen | 25µg/kg | 6 | Post | Rat, contusion | Decreased | (Majidpoor et al., 2020) |
| VEGF | Angiogenesis, vasculogenesis, and endothelial cell growth | Premarin | 100µg/kg | 8 | Post | Rat, contusion | Increased | (Haque et al., 2022) |
|  |  |  | 1mg/kg | Implant | Post | Rat, compression | Increased | (Chen et al., 2010) |
| Zo-1* | Scaffolding; links tight junction transmembrane proteins | Estrogen | 300µg/kg | 3 | Post | Rat, contusion | Increased | (Lee et al., 2015) |

* denotes proteins used in the STRING analysis. Green indicates a beneficial effect, orange indicates a temporarily or some beneficial effect, red indicates no effect.

Abbreviations: Abcc8: ATP binding cassette subfamily C member 8; Aim2: absent in melanoma 2; Akt: serine/threonine kinase 1; Ang1: angiopoietin 1; Ang-2: angiotensin II; APC: APC regulator of WNT signalling pathway; ASC: Apoptosis-associated speck-like protein containing a caspase recruitment domain; Atg5: autophagy related 5; Atg7: autophagy related 7; AQP4: aquaporin 4; BAD: Bcl2-associated agonist of cell death; BAX: Bcl2-associated X apoptosis regulator; Bcl-2: Bcl-2 apoptosis regulator; Bcl-Xl: BCL2 like 1; BDNF: brain-derived neurotrophic factor; Cd11: cluster of differentiation 11; Cd11b: cluster of differentiation 11β; Cd13: cluster of differentiation 13; Cd31: cluster of differentiation 31; Cd61: cluster of differentiation 61; Cd68: cluster of differentiation 68; Cd163: cluster of differentiation 163; CGRP: calcitonin gene-related peptide; c-Jun: jun proto-oncogene; CNPase: cyclic nucleotide 3’-phosphodiesterase; COX-2: cytochrome c oxidase II; CREB: cAMP responsive element binding protein; ERK: extracellular signal-regulated kinase; Flk-1: kinase insert domain receptor; Flt-1: Fms related receptor tyrosine kinase 1; Gap43: growth associated protein 43; GDNF: glial cell derived neurotrophic factor; GFAP: glial fibrillary acidic protein; Glt-1: glutamate/aspartate transporter II; Glut-1: glucose transporter type 1; GROα: growth-regulated α protein; GSTπ: glutathione S-transferase π; Iba1: ionized calcium-binding adapter molecule 1; IκBα: NF-κB inhibitor α; IL-1α: interleukin 1α; IL-1β: interleukin 1β; IL-6: interleukin 6; IL-18: interleukin 18; iNos: inducible nitric oxide synthase; JNK1/2: mitogen-activated protein kinase 8; Kir4.1: potassium inwardly rectifying channel subfamily J member 10; Lc3: microtubule associated protein 1 light chain 3; MAG: myelin associated glycoprotein; MBP: myelin basic protein; MCP-1: monocyte chemoattractant protein-1; Mip-1α: macrophage inflammatory protein 1α; Mip-1β: macrophage inflammatory protein 1β; Mip-2α: macrophage inflammatory protein 2α; miR-106a: microRNA 106a; miR-17: microRNA 17; miR-206: microRNA 206; miR-21: microRNA 21; miR-7-1: microRNA 7-1; Mmp1: matrix metallopeptidase 1; Mmp2: matrix metallopeptidase 2; Mmp3: matrix metallopeptidase 3; Mmp9: matrix metallopeptidase 9; Mmp10: matrix metallopeptidase 10; Mmp13: matrix metallopeptidase 13; MOG: myelin oligodendrocyte glycoprotein; MPO: myeloperoxidase; NeuN: neuronal nuclei; NF-H: neurofilament heavy chain; NF-κB: nuclear factor κB; NFP: serine/threonine receptor-like kinase NFP; NG2: neuronal-glial antigen 2; Nrlc4: NLR family CARD domain containing 4; Nlrp1: NACHT, LRR, and PYD domains-containing protein 1; Nlrp1b: NACHT, LRR, and PYD domains-containing protein 1b; Nlrp3: NACHT, LRR, and PYD domains-containing protein 3; NogoA: reticulon-4; Nox2: NADPH oxidase isoform 2; O4: forehead box protein O4; Olig2: oligodendrocyte transcription factor 2; OMgp: oligodendrocyte-myelin glycoprotein; PI3K: phosphatidylinositol 3-kinase; PTEN: phosphatidylinositol 3,4,5-triphosphate 3-phosphatase and dual-specificity protein phosphatase; RhoA: ras homolog family member A; Src: proto-oncogene tyrosine-protein kinase; Sqstm1: sequestosome-1; Tie2: tyrosine-protein kinase receptor 2; Tlr4: Toll-like receptor 4; TNF-α: tumor necrosis factor α; Trpm4: transient receptor potential cation channel subfamily M member 4; Txnip: thioredoxin-interaction protein; VEGF: vascular endothelial growth factor; Zo-1: tight protein junction 1
